# R-Spondin Mimetic, SZN-043, Induced Proliferation and Expression of Wnt Target Genes, Two Impaired Features in Human Alcohol-Associated Liver Disease

**DOI:** 10.1101/2024.11.19.624277

**Authors:** Trevor Fisher, Mehaben Patel, Shalaka Deshmukh, Darshini Shah, Chenggang Lu, Maureen Newman, Jay Ye, Russell Fletcher, Geertrui F. Vanhove, Jay Tibbitts, Yang Li, Nicholas J Skill, Zhihong Yang, Suthat Liangpunsakul, Helene Baribault

## Abstract

Liver regeneration is impaired in patients suffering from alcohol-associated liver (ALD) diseases. Wnt ligands and their FZD receptors are dysregulated in diseased livers. R-spondin and their receptors are known to regulate Wnt activity via the stabilization of FZD receptors. Here, we investigated the components of the Wnt and R-Spondin-signaling pathways and their activity in patients with ALD. We found that while hepatocytes retained high levels of differentiation markers such as *ASGR1* and *ASGR2*, the expression of two R-spondin co-receptors, *LGR4* and *LGR5*, and of Wnt target genes, *CYP1A2* and others, were strongly reduced.

SZN-043, a hepatocyte-targeted R-Spondin mimetic, is a new investigational drug that stimulates the physiological Wnt repair pathway and proliferation of hepatocytes. Here, we show that SZN-043 induced hepatocyte proliferation in all models tested, including humanized mouse livers, a chronic-binge alcohol-induced liver injury, and a CCl_4_-induced fibrosis mouse model. Altogether, SZN-043 could be beneficial for the treatment of ALD.

## INTRODUCTION

Alcohol-associated liver disease (ALD) manifests through a spectrum of stages, each representing varying degrees of liver damage (1). Progression typically begins with alcohol-associated steatosis, characterized by the accumulation of fat in hepatocytes due to chronic alcohol consumption (2). The disease can advance to alcohol-associated hepatitis (AH), a clinical syndrome marked by inflammation and hepatocellular injury (2). AH presents with symptoms such as jaundice and is associated with an elevated risk of liver-related complications (3). Importantly, AH can occur at any stage of ALD progression. In fact, up to 80% of patients diagnosed with severe AH (sAH), as indicated by a high model for end-stage liver disease (MELD) score exceeding 20, may have underlying cirrhosfis (3). Mild to moderate cases of AH often follow a self-limiting trajectory, resolving without significant intervention (4). However, sAH poses a considerable risk, with markedly higher mortality rates (5). In fact, approximately 15 to 20% of patients diagnosed with sAH may succumb to the condition within just one month, with this figure rising to around 30% at the three-month mark (6). The liver is known for its ability to regenerate. Yet, its capacity for regrowth is severely compromised as liver disease progresses (for review (7)). Studies have consistently demonstrated a correlation between enhanced hepatocyte proliferative capacity and improved survival outcomes in patients with sAH (8–10). Thus, interventions aimed at promoting hepatocyte proliferation may hold promise for mitigating the severity of AH and enhancing patient survival or reducing the need for liver transplantation.

The Wnt/β-catenin signaling pathway serves as a fundamental regulator of liver tissue homeostasis, orchestrating various cellular processes crucial for liver function and repair, especially in response to injury (11, 12). Among its pivotal roles, this signaling pathway governs hepatocyte proliferation, regeneration, and maturation, exerting direct control over the expression of key cell cycle regulators like Cyclin D1 (12–19). Furthermore, Wnt/β-catenin signaling serves as a master regulator of liver zonation, a process crucial for defining the metabolic identities of hepatocytes within distinct regions of the liver lobule (20, 21). The intricate involvement of Wnt/β-catenin signaling in hepatocyte function, proliferation, and zonation highlights its significance in liver biology and pathology (22). Insights into the mechanisms governing this pathway hold promise for advancing our understanding of liver diseases and may pave the way for innovative therapeutic approaches aimed at promoting liver regeneration and mitigating liver injury.

R-spondins (RSPOs), which are known for their ability to amplify Wnt signaling, play crucial roles in liver biology, particularly in liver zonation and regeneration (23–25). RSPOs enhance Wnt signaling, by stabilizing the Frizzled (FZD) and LRP co-receptors of Wnt proteins at the plasma membrane, thereby influencing various cellular processes (26). In the liver, RSPOs have been shown to promote regeneration and maintain zonation, especially in response to injury induced by agents such as carbon tetrachloride (CCl_4_) or hepatectomy (23–25). However, the effects of RSPOs extend beyond hepatocytes, as they can elicit responses in other tissues, including small intestinal hyperplasia (27, 28). While enhancing Wnt signaling through RSPOs may be beneficial for liver regeneration, caution is warranted due to the potential downside associated with excessive Wnt signaling in hepatic stellate cells (29). This duality underscores the need for a nuanced understanding of Wnt signaling modulation in the context of liver regeneration and fibrosis.

The development of a hepatocyte-targeted R-spondin mimetic, such as SZN-043, represents a promising approach to address the lack of specificity associated with systemically administered RSPOs (28). By specifically targeting hepatocytes by binding to the ubiquitin E3 ligases ZNRF3 and RNF43, as well as the hepatocyte-specific receptor ASGR1, SZN-043 offers a more precise and targeted therapeutic strategy for promoting liver regeneration and function (28). Previous studies have demonstrated the ability of a precursor of SZN-043, αASGR1-RSPO2-RA-IgG, to upregulate Wnt target genes and promote hepatocyte proliferation in the liver without affecting the intestine (28). SZN-043, a version optimized for pharmacological properties such as stability and specificity while retaining its binding affinity and efficacy in promoting hepatocyte proliferation, holds significant potential as a therapeutic intervention for liver diseases, offering a targeted and effective pharmacological approach to promote liver regeneration and function while minimizing off-target effects.

In this study, our primary objective was to investigate the role of the Wnt signaling pathway in patients with severe ALD and to explore the therapeutic potential of SZN-043, a hepatocyte-targeted R-spondin mimetic, in various preclinical models of ALD. We conducted a transcriptomic analysis of liver tissue obtained from patients diagnosed with alcohol-associated cirrhosis in two independent cohorts (**Supplementary Tables S1 and S2**). Our goal was to assess the expression levels of Wnt target genes in the liver of these patients. We observed a significant reduction in Wnt target gene expression, accompanied by a lack of hepatocyte proliferation. Furthermore, we investigated the expression of key genes involved in the mechanism of action of SZN-043, including *ASGR1*, *ZNRF3*, and *RNF43* and observed that the expression levels of these genes remained unaffected in patients with ALD, indicating that the molecular targets of SZN-043 were still present and potentially functional in the context of severe ALD. This finding provides a rationale for exploring the therapeutic effects of SZN-043 in preclinical models of ALD, with the aim of restoring Wnt signaling and promoting hepatocyte proliferation as a potential treatment strategy for severe ALD. First, we investigated the impact of SZN-043 on hepatocyte expansion and liver function in healthy mice. Additionally, we examined its effects on human hepatocytes transplanted into FRG® KO (FRG) mice with humanized livers. These experiments aimed to assess the safety and efficacy of SZN-043 in promoting hepatocyte proliferation in healthy and humanized liver environments. Furthermore, we evaluated the therapeutic potential of SZN-043 in commonly used alcohol-induced liver injury models. We also assessed the effects of SZN-043 on hepatocyte expansion in a CCl_4_-induced fibrosis model, considering the relevance of fibrosis in the context of ALD progression (30, 31). Our results demonstrated that SZN-043 effectively increased Wnt signaling and promoted hepatocyte proliferation in alcohol-induced liver injury models. Moreover, we observed that SZN-043 induced hepatocyte proliferation and reduced fibrosis in the CCl_4_-induced fibrosis model. These findings suggest that SZN-043 holds promise as a potential treatment for alcohol-associated hepatitis, offering a novel therapeutic approach to address this challenging condition by targeting the dysregulated Wnt signaling pathway and promoting hepatocyte regeneration while mitigating fibrosis.

## METHODS

Methods can be found in the supplemental section.

## RESULTS

### Dysregulation of Wnt signaling pathway components in the liver of patients with alcohol-associated cirrhosis

Our investigation into the expression patterns of Wnt ligands, R-spondin modulators, and Wnt receptors in the liver of patients with alcohol-associated cirrhosis yielded some intriguing findings (**Figure 1**). Notably, we observed significant elevations in the expression of multiple Wnt ligands, such as *WNT2B, WNT4, WNT5B, WNT7B, WNT9A, WNT10A*, and *WNT10B*, in the liver of patients with alcohol-associated cirrhosis, as indicated by both RT-qPCR **(Figure 1A)** and RNA-seq analysis **(Table 1)**.

**Figure 1.**
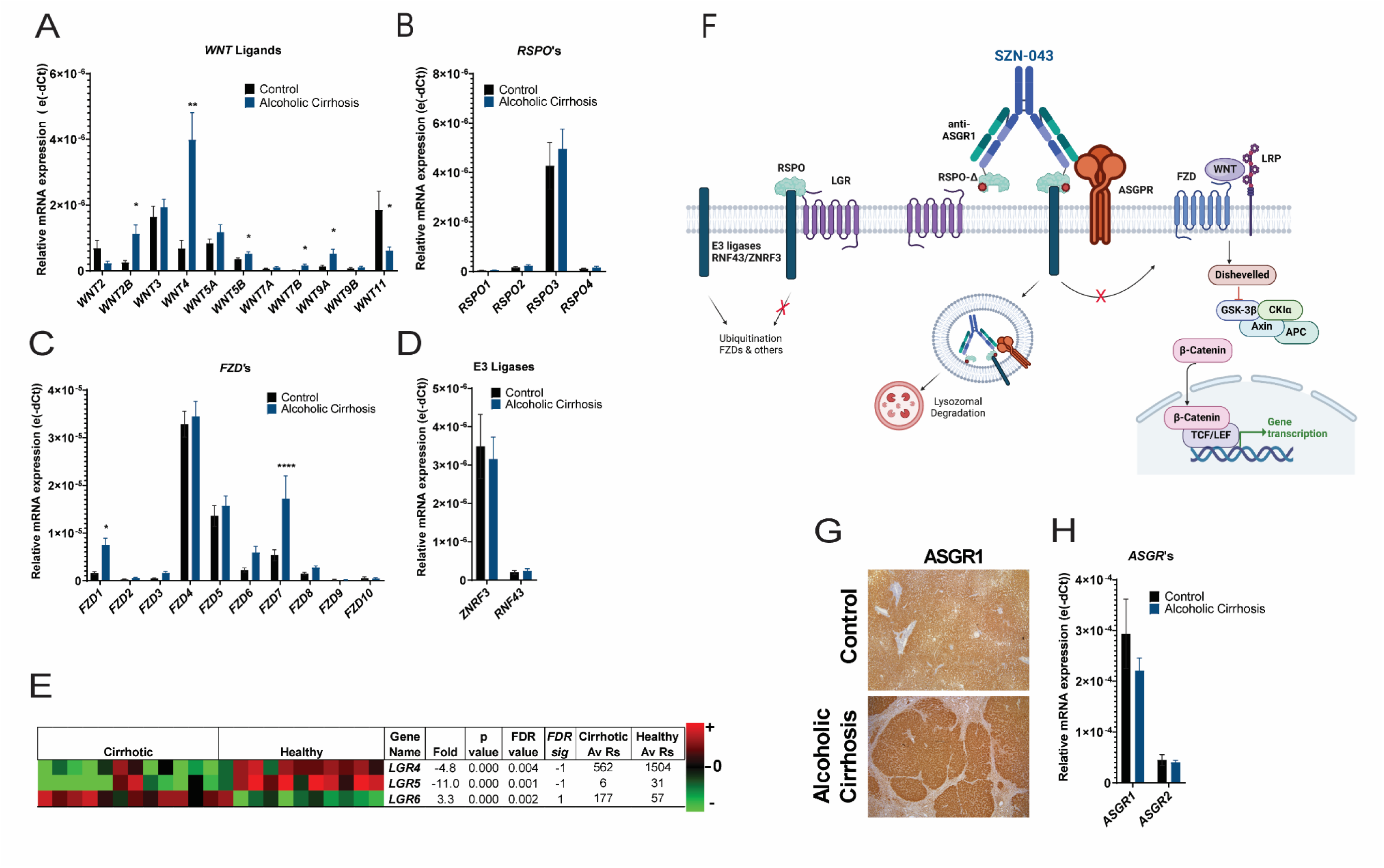
Expression of Wnt Signaling Pathway Components in Alcohol-Associated Cirrhotic and Control Livers from Human Patients. (A) QPCR expression analysis of 11 *WNT*, (B) 4 *RSPO* and (C) 10 *FZD* family members, (D) QPCR expression analysis of *RNF43*, *ZNRF3* (H) *ASGR1* and *ASGR2* in control and alcohol-associated cirrhosis samples. Results are presented as (e(dCt)) to reflect the relative abundance of different family members. (E) Bulk RNAseq expression analysis of 3 *LGR4*, *LGR5* and *LGR6* in control and alcohol-associated cirrhosis samples. FDR is the false discovery rate, FDR sig = -1 means that there is a significant reduction in alcohol-associated cirrhosis samples using FDR < 0.05, FDR sig = 1 means that there is a significant increase in alcohol-associated cirrhosis samples using FDR < 0.05, Av Rs is the average number of readings (F) Schematic representation of the mechanism by which the RSPO mimetic, SZN-043, induces hepatocyte-targeted Wnt signaling (created with BioRender.com). (From left to right) In absence of RSPO, the E3 ligases, RNF43 and ZNRF3, induce the ubiquitination of FZD receptors and subsequent degradation. Upon binding of RSPO to the E3 ligases and its co-receptor LGR, RSPO prevents FZD ubiquitination and thereby induces the stabilization of FZD receptors. SZN-043 is a fusion antibody-based ligand containing a mutated RSPO which does not bind the LGR receptor, but still binds E3 ligases and an anti-ASGR1 binding domain. Upon binding to ASGR1, expressed predominantly in hepatocytes, and E3 ligases, SZN-043 induces Wnt activity, via the stabilization of its receptors. SZN-043 may be internalized along with ASGR1 into lysosomes. (G) Histoimmunochemistry of control and alcohol-associated cirrhosis samples liver sections using anti-human ASGR1 antibodies.

**Table 1.**
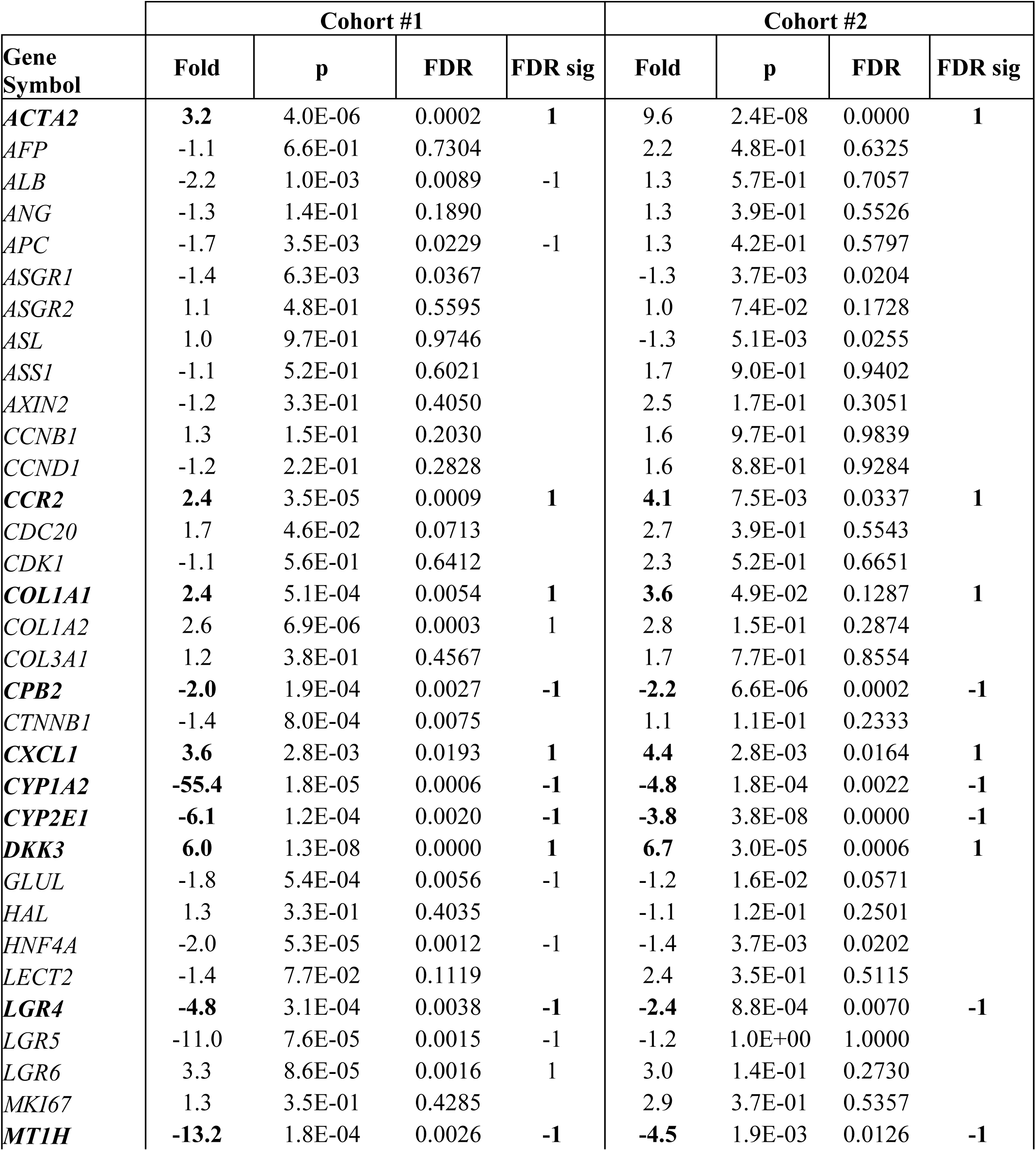
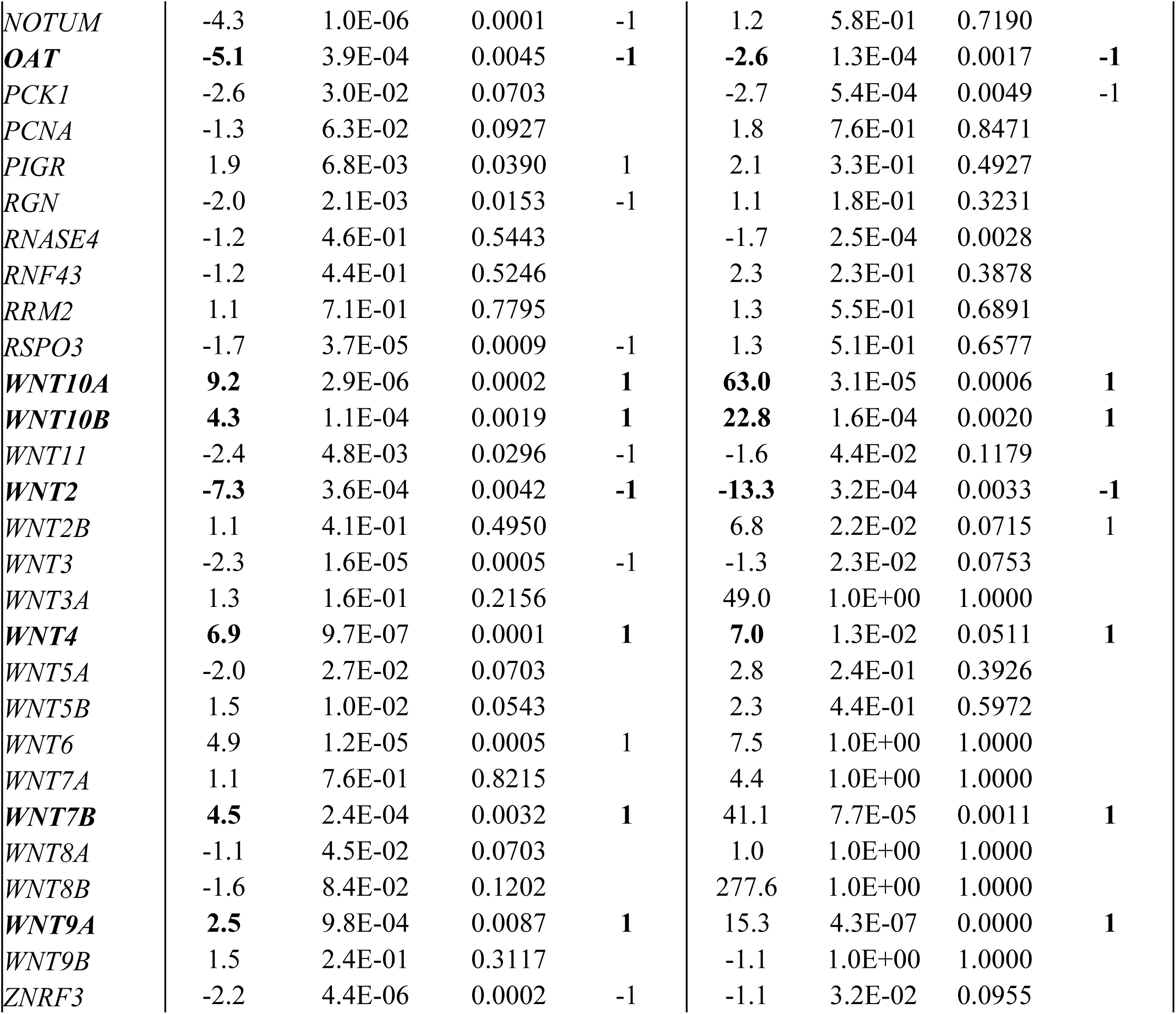
Selected genes comparative RNAseq analysis in two cohorts of control and alcoholic cirrhosis liver samples. FDR: false discovery rate, FDRsig = 1, significant fold increase when using FDR < 0.05, FDRsig = -1, significant fold decrease when using FDR < 0.05. Genes that were significantly elevated or reduced in both cohorts are highlighted in bold.

| Gene Symbol | Cohort #1 |  |  |  | Cohort #2 |  |  |  |
| --- | --- | --- | --- | --- | --- | --- | --- | --- |
|  | Fold | p | FDR | FDR sig | Fold | p | FDR | FDR sig |
| **ACTA2** | **3.2** | 4.0E-06 | 0.0002 | **1** | 9.6 | 2.4E-08 | 0.0000 | **1** |
| *AFP* | -1.1 | 6.6E-01 | 0.7304 |  | 2.2 | 4.8E-01 | 0.6325 |  |
| *ALB* | -2.2 | 1.0E-03 | 0.0089 | -1 | 1.3 | 5.7E-01 | 0.7057 |  |
| *ANG* | -1.3 | 1.4E-01 | 0.1890 |  | 1.3 | 3.9E-01 | 0.5526 |  |
| *APC* | -1.7 | 3.5E-03 | 0.0229 | -1 | 1.3 | 4.2E-01 | 0.5797 |  |
| *ASGR1* | -1.4 | 6.3E-03 | 0.0367 |  | -1.3 | 3.7E-03 | 0.0204 |  |
| *ASGR2* | 1.1 | 4.8E-01 | 0.5595 |  | 1.0 | 7.4E-02 | 0.1728 |  |
| *ASL* | 1.0 | 9.7E-01 | 0.9746 |  | -1.3 | 5.1E-03 | 0.0255 |  |
| *ASS1* | -1.1 | 5.2E-01 | 0.6021 |  | 1.7 | 9.0E-01 | 0.9402 |  |
| *AXIN2* | -1.2 | 3.3E-01 | 0.4050 |  | 2.5 | 1.7E-01 | 0.3051 |  |
| *CCNB1* | 1.3 | 1.5E-01 | 0.2030 |  | 1.6 | 9.7E-01 | 0.9839 |  |
| *CCND1* | -1.2 | 2.2E-01 | 0.2828 |  | 1.6 | 8.8E-01 | 0.9284 |  |
| **CCR2** | **2.4** | 3.5E-05 | 0.0009 | **1** | **4.1** | 7.5E-03 | 0.0337 | **1** |
| *CDC20* | 1.7 | 4.6E-02 | 0.0713 |  | 2.7 | 3.9E-01 | 0.5543 |  |
| *CDK1* | -1.1 | 5.6E-01 | 0.6412 |  | 2.3 | 5.2E-01 | 0.6651 |  |
| **COL1A1** | **2.4** | 5.1E-04 | 0.0054 | **1** | **3.6** | 4.9E-02 | 0.1287 | **1** |
| *COL1A2* | 2.6 | 6.9E-06 | 0.0003 | 1 | 2.8 | 1.5E-01 | 0.2874 |  |
| *COL3A1* | 1.2 | 3.8E-01 | 0.4567 |  | 1.7 | 7.7E-01 | 0.8554 |  |
| **CPB2** | **-2.0** | 1.9E-04 | 0.0027 | **-1** | **-2.2** | 6.6E-06 | 0.0002 | **-1** |
| *CTNNB1* | -1.4 | 8.0E-04 | 0.0075 |  | 1.1 | 1.1E-01 | 0.2333 |  |
| **CXCL1** | **3.6** | 2.8E-03 | 0.0193 | **1** | **4.4** | 2.8E-03 | 0.0164 | **1** |
| **CYP1A2** | **-55.4** | 1.8E-05 | 0.0006 | **-1** | **-4.8** | 1.8E-04 | 0.0022 | **-1** |
| **CYP2E1** | **-6.1** | 1.2E-04 | 0.0020 | **-1** | **-3.8** | 3.8E-08 | 0.0000 | **-1** |
| **DKK3** | **6.0** | 1.3E-08 | 0.0000 | **1** | **6.7** | 3.0E-05 | 0.0006 | **1** |
| *GLUL* | -1.8 | 5.4E-04 | 0.0056 | -1 | -1.2 | 1.6E-02 | 0.0571 |  |
| *HAL* | 1.3 | 3.3E-01 | 0.4035 |  | -1.1 | 1.2E-01 | 0.2501 |  |
| *HNF4A* | -2.0 | 5.3E-05 | 0.0012 | -1 | -1.4 | 3.7E-03 | 0.0202 |  |
| *LECT2* | -1.4 | 7.7E-02 | 0.1119 |  | 2.4 | 3.5E-01 | 0.5115 |  |
| **LGR4** | **-4.8** | 3.1E-04 | 0.0038 | **-1** | **-2.4** | 8.8E-04 | 0.0070 | **-1** |
| *LGR5* | -11.0 | 7.6E-05 | 0.0015 | -1 | -1.2 | 1.0E+00 | 1.0000 |  |
| *LGR6* | 3.3 | 8.6E-05 | 0.0016 | 1 | 3.0 | 1.4E-01 | 0.2730 |  |
| *MKI67* | 1.3 | 3.5E-01 | 0.4285 |  | 2.9 | 3.7E-01 | 0.5357 |  |
| **MT1H** | **-13.2** | 1.8E-04 | 0.0026 | **-1** | **-4.5** | 1.9E-03 | 0.0126 | **-1** |
| *NOTUM* | -4.3 | 1.0E-06 | 0.0001 | -1 | 1.2 | 5.8E-01 | 0.7190 |  |
| ***OAT*** | **-5.1** | 3.9E-04 | 0.0045 | **-1** | **-2.6** | 1.3E-04 | 0.0017 | **-1** |
| *PCK1* | -2.6 | 3.0E-02 | 0.0703 |  | -2.7 | 5.4E-04 | 0.0049 | -1 |
| *PCNA* | -1.3 | 6.3E-02 | 0.0927 |  | 1.8 | 7.6E-01 | 0.8471 |  |
| *PIGR* | 1.9 | 6.8E-03 | 0.0390 | 1 | 2.1 | 3.3E-01 | 0.4927 |  |
| *RGN* | -2.0 | 2.1E-03 | 0.0153 | -1 | 1.1 | 1.8E-01 | 0.3231 |  |
| *RNASE4* | -1.2 | 4.6E-01 | 0.5443 |  | -1.7 | 2.5E-04 | 0.0028 |  |
| *RNF43* | -1.2 | 4.4E-01 | 0.5246 |  | 2.3 | 2.3E-01 | 0.3878 |  |
| *RRM2* | 1.1 | 7.1E-01 | 0.7795 |  | 1.3 | 5.5E-01 | 0.6891 |  |
| *RSPO3* | -1.7 | 3.7E-05 | 0.0009 | -1 | 1.3 | 5.1E-01 | 0.6577 |  |
| ***WNT10A*** | **9.2** | 2.9E-06 | 0.0002 | **1** | **63.0** | 3.1E-05 | 0.0006 | **1** |
| ***WNT10B*** | **4.3** | 1.1E-04 | 0.0019 | **1** | **22.8** | 1.6E-04 | 0.0020 | **1** |
| *WNT11* | -2.4 | 4.8E-03 | 0.0296 | -1 | -1.6 | 4.4E-02 | 0.1179 |  |
| ***WNT2*** | **-7.3** | 3.6E-04 | 0.0042 | **-1** | **-13.3** | 3.2E-04 | 0.0033 | **-1** |
| *WNT2B* | 1.1 | 4.1E-01 | 0.4950 |  | 6.8 | 2.2E-02 | 0.0715 | 1 |
| *WNT3* | -2.3 | 1.6E-05 | 0.0005 | -1 | -1.3 | 2.3E-02 | 0.0753 |  |
| *WNT3A* | 1.3 | 1.6E-01 | 0.2156 |  | 49.0 | 1.0E+00 | 1.0000 |  |
| ***WNT4*** | **6.9** | 9.7E-07 | 0.0001 | **1** | **7.0** | 1.3E-02 | 0.0511 | **1** |
| *WNT5A* | -2.0 | 2.7E-02 | 0.0703 |  | 2.8 | 2.4E-01 | 0.3926 |  |
| *WNT5B* | 1.5 | 1.0E-02 | 0.0543 |  | 2.3 | 4.4E-01 | 0.5972 |  |
| *WNT6* | 4.9 | 1.2E-05 | 0.0005 | 1 | 7.5 | 1.0E+00 | 1.0000 |  |
| *WNT7A* | 1.1 | 7.6E-01 | 0.8215 |  | 4.4 | 1.0E+00 | 1.0000 |  |
| ***WNT7B*** | **4.5** | 2.4E-04 | 0.0032 | **1** | 41.1 | 7.7E-05 | 0.0011 | **1** |
| *WNT8A* | -1.1 | 4.5E-02 | 0.0703 |  | 1.0 | 1.0E+00 | 1.0000 |  |
| *WNT8B* | -1.6 | 8.4E-02 | 0.1202 |  | 277.6 | 1.0E+00 | 1.0000 |  |
| ***WNT9A*** | **2.5** | 9.8E-04 | 0.0087 | **1** | 15.3 | 4.3E-07 | 0.0000 | **1** |
| *WNT9B* | 1.5 | 2.4E-01 | 0.3117 |  | -1.1 | 1.0E+00 | 1.0000 |  |
| *ZNRF3* | -2.2 | 4.4E-06 | 0.0002 | -1 | -1.1 | 3.2E-02 | 0.0955 |  |

Conversely, *WNT2* and *WNT11* exhibited a trend toward reduction **(Figure 1A and Table 1)**. While other Wnt ligands showed small and inconsistent changes across the datasets analyzed, these findings collectively suggest dysregulation of Wnt ligand expression in the liver of patients with alcohol-associated cirrhosis. Regarding Wnt receptors, the expression of *FZD4* and *FZD5*, which are prominently expressed in human hepatocytes, did not display significant differences between healthy and cirrhotic livers **(Figure 1B)**. However, *FZD1* and *FZD7*, expressed at lower levels in control livers, were found to be elevated in cirrhotic samples. This observation suggests a potential alteration in Wnt receptor expression profiles associated with cirrhosis, which may have implications for Wnt signaling dynamics and liver pathology.

The expression of R-spondins, which enhances Wnt signaling by inhibiting the activity of E3 ligase receptors (ZNRF3 and RNF43), thereby prevents degradation of FZDs and LRPs (25), yielded interesting results. We found no significant differences in the expression of *RSPO3* or other R-spondins between cirrhotic and control livers (**Figure 1C**), indicating that the levels of these Wnt signaling enhancers remained stable in the context of liver cirrhosis. Similarly, there were no discernible differences in the expression of *ZNRF3* or *RNF43*, the E3 ligase receptors targeted by R-spondins, suggesting that the degradation mechanisms for FZDs and LRPs may not be significantly altered in cirrhotic livers. However, our analysis of RNA-seq data revealed a notable downregulation of the R-spondin co-receptor genes *LGR4* and *LGR5* in both cohorts studied (**Table 1**). Since *LGR4* is the most abundant member of the LGR family expressed in hepatocytes, this decrease in expression could potentially impair Wnt activity in these cells (**Figure 1F**), thus affecting Wnt signaling dynamics in the context of liver cirrhosis. Overall, our findings shed light on alterations in the expression of Wnt ligands, receptors, and modulators in cirrhotic livers, indicating dysregulation of the Wnt signaling pathway in the context of liver cirrhosis. These observations provide valuable insights into the molecular mechanisms underlying the pathogenesis of ALD, contributing to our understanding of this complex liver disease.

### ASGR1, used for SZN-043 hepatocyte targeting, is abundantly expressed in hepatocytes of patients with alcohol-associated cirrhosis

SZN-043, an antibody-based fusion protein designed for targeted Wnt signaling modulation in hepatocytes, consists of a binding domain specific to the hepatocyte-specific ASGR1 receptor and a mutant R-spondin capable of binding to ZNRF3 and RNF43 (**Figure 1F**) but unable to interact with LGR receptors(28). This design enabled SZN-043 to selectively stimulate Wnt signaling in hepatocytes, mimicking the effect of R-spondins. To ensure the efficacy of SZN-043 in cirrhotic livers, the expression of the critical SZN-043 targeting receptor ASGR1 in hepatocytes was evaluated. This analysis involved comparing the expression levels of ASGR1 and its co-receptor ASGR2 in cirrhotic and healthy livers using both immunostaining (**Figure 1G**) and RT-qPCR (**Figure 1H**). Our findings demonstrated that the levels of ASGR1 expression, as measured by RT-qPCR, were similar between normal and cirrhotic livers. Additionally, immunohistochemistry staining revealed abundant ASGR1 expression in cirrhotic livers, confirming the preservation of this key targeting protein in the context of alcohol-induced liver injury.

### Reduction in the expression of Wnt target genes in patients with alcohol-associated cirrhosis

Global transcriptome profiling by RNA-Seq revealed a significant reduction in the expression of several Wnt/β-catenin target genes in the livers of patients with alcohol-associated cirrhosis. Among these genes, *CYP1A2*, a well-known Wnt/β-catenin target gene responsible for metabolizing xenobiotics and predominantly expressed in the pericentral zone (32), exhibited the most substantial downregulation. In cohort #1, *CYP1A2* expression was reduced by 55.4-fold compared to healthy controls (**Table 1**). This reduction in *CYP1A2* expression was further confirmed in a second independent alcohol-associated cirrhosis cohort (cohort #2), with a 4.8-fold decrease observed (**Table 1**). RT-qPCR analysis also validated the significant reduction in *CYP1A2* expression (**Figure 2A**). Immunostaining analysis further corroborated these findings, showing a decrease in CYP1A2 protein levels in cirrhotic liver samples (**Figure 2B**). In addition to *CYP1A2*, the expression of other Wnt/β-catenin target genes, such as *CYP2E1, GLUL, LGR5*, *OAT* and *ZNRF3*, was also reduced in the livers of cirrhotic patients, as evidenced by RNA-Seq analysis (**Table 1**). These findings underscore the dysregulation of Wnt/β-catenin signaling in alcohol-associated cirrhosis and its downstream effects on the expression of genes involved in xenobiotic metabolism and liver function.

**Figure 2.**
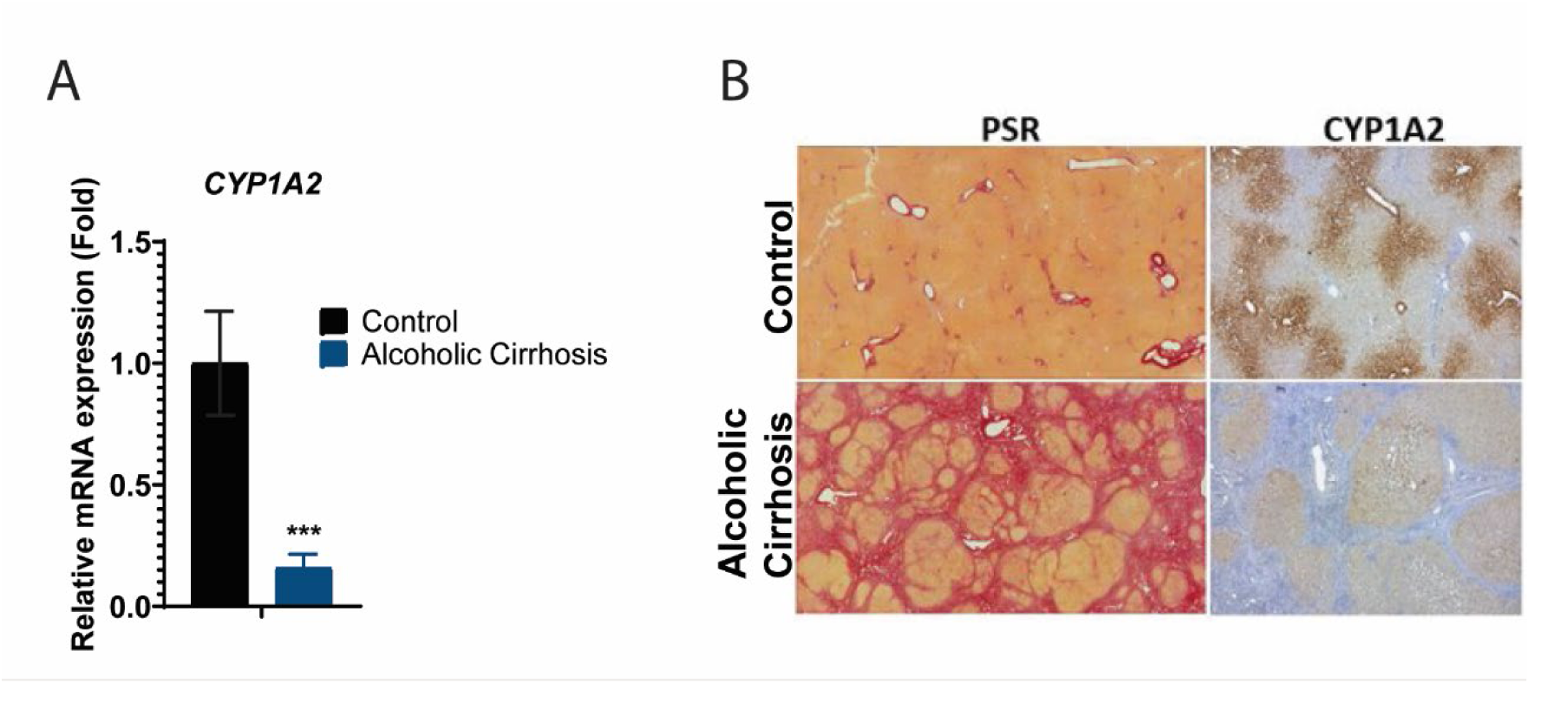
Expression of the Wnt target gene, CYP1A2, and (PSR) Staining in Alcohol-Associated Cirrhotic and Control Livers from Human Patients. (A) QPCR expression analysis of *CYP1A2*. (B) PSR histological staining and CYP1A2 immunostaining of control and alcohol-associated cirrhosis liver sections.

The expression of other Wnt target genes, such as *AXIN2*, *CCND1* and *NOTUM* were not consistently and significantly changed. It should be noted that unlike the hepatocyte-enriched or –specific target genes mentioned above, *AXIN2*, *CCND1* and *NOTUM* are highly expressed in mesenchymal cells also. Given that SZN-043 is not active on mesenchymal cells, this discrepancy could be best explained by the large increase in mesenchymal cells in ALD tissue samples, thereby confounding the results.

### Lack of spontaneous regeneration markers in the livers of patients with alcohol-associated cirrhosis

The Wnt/β-catenin pathway plays a crucial role in liver regeneration, particularly in response to liver injury (33). It has been observed that improved hepatocyte proliferation can lead to better outcomes and increased survival in patients with sAH (8, 9). However, the reduced expression of Wnt target genes, as observed in alcohol-associated cirrhotic livers, suggests decreased activity of this pathway, which may contribute to insufficient hepatocyte proliferation. To further investigate whether hepatocyte regeneration was compromised at the molecular level, we examined the expression of genes involved in cell cycle regulation (**Table 1** and **Figure S1**). These genes control distinct phases of the cell cycle, including the G1/S transition, S phase, and G2/M transition, which are critical for cell proliferation (34, 35).

Specifically, we looked at the expression of *PCNA* and *CCND1*, which regulate the G1/S phase transition; *RRM2*, which is involved in DNA synthesis during the S phase; and *CCNB1*, *MKI67*, *CDC20*, and *CDK1*, which regulate entry into the G2/M phase (34, 35). Interestingly, none of these genes were significantly elevated in cirrhotic livers compared to healthy control livers in either cohort (**Table 1**). Consistent with these results, double immunofluorescence staining of these tissue samples with the proliferation marker, Ki-67, and the hepatocyte-specific marker, HNF4A, showed few positive cells for both markers, in either the healthy or cirrhotic tissue sections (**Figure S1**). This suggests that spontaneous hepatocyte proliferation, as indicated by the upregulation of cell cycle-related genes, does not occur in response to cirrhotic injuries in liver tissues of patients with ALD. This finding further supports the notion of impaired hepatocyte regeneration in alcohol-associated cirrhosis, potentially due to reduced Wnt/β-catenin pathway activity.

### SZN-043 induced Wnt target genes and proliferation markers in a dose-dependent manner in mice

Investigations into potential treatments for ALD have led to the consideration of molecules that can stimulate hepatocyte proliferation and restore the expression of Wnt target genes, given the observed reduction in these genes in cirrhotic livers. SZN-043, a compound previously shown to induce hepatocyte-targeted proliferation and increase Wnt target gene expression in mouse livers (28), presents an opportunity for ALD treatment.

We evaluated the single-dose pharmacokinetic (PK) profile of SZN-043 and its effects on selected pharmacodynamic (PD) biomarkers, following intravenous (IV) bolus injection in C57BL/6J (B6) mice. Serum concentrations of SZN-043 increased with dose, but not in a dose-dependent manner in the dose range of 0.3 mg/kg to 200 mg/kg (**Figure 3A**). The clearance and mean residence time of SZN-043 significantly decreased with increasing doses (**Table 2**). These findings are consistent with ASGR being a high-capacity and rapidly internalizing receptor, abundantly and specifically expressed on the sinusoidal surface of hepatocytes, which internalizes ligands via clathrin-dependent receptor-mediated endocytosis, followed by degradation of the ligands in the lysosome (36). The elimination of SZN-043 most likely occurs primarily by target-mediated drug disposition (TMDD) associated with ASGR-mediated processes (37).

**Figure 3.**
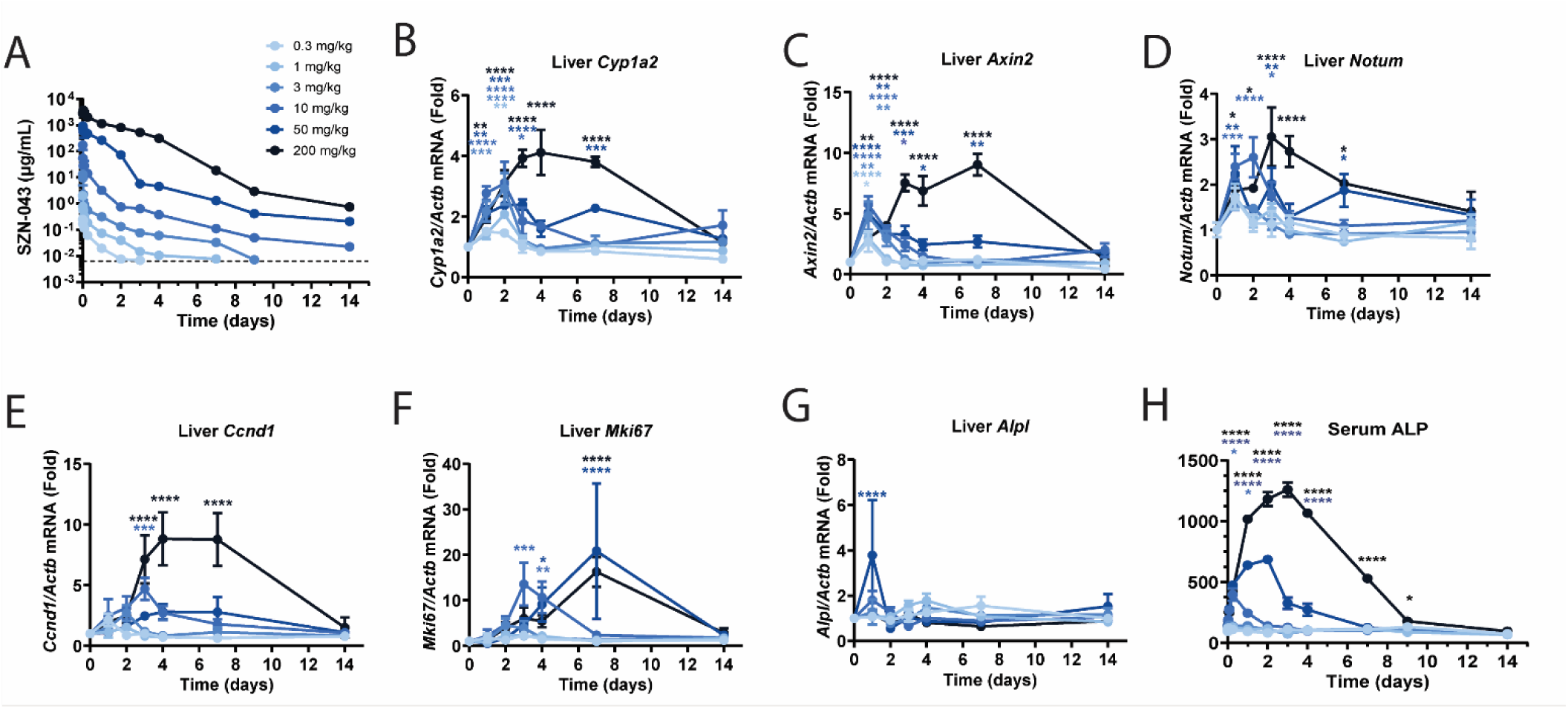
Dose Response Analysis of SZN-043 effect on the Expression of Wnt Target Genes, its Feedback Inhibitors and Proliferation Markers in Mice. (A) Serum exposure of SZN-043 over time after a single intravenous administration of various SZN-043 doses ranging from 0.3 to 200 mg/kg. (B-G) Time course qPCR expression analysis of (B) *Cyp1a2*, (C) *Axin2*, (D) *Notum*, (E) *Ccnd1*, (F) *Mki67*and (G) *Alpl* and (H) serum ALP level.

**Table 2.** Pharmacokinetic Parameters of SZN-043 in Mice Following a Single IV Bolus Dose. AUC_inf_, area under the serum concentration curve from time 0 to time infinity; C_0,_ estimated serum concentration at time zero; CL, clearance; MRT, mean residence time; Terminal t_1/2_, terminal half-life; Vc, central compartment volume of distribution.

| **Dose**<br>**(mg/kg)** | **C<sub>0</sub>**<br>**(µg/mL)** | **AUC<sub>inf</sub>**<br>**(µg-day/mL)** | **V<sub>c</sub>**<br>**(mL/kg)** | **CL**<br>**(mL/day/kg)** | **Terminal t<sub>1/2</sub>**<br>**(days)** | **MRT**<br>**(days)** |
| --- | --- | --- | --- | --- | --- | --- |
| 0.3 | 0.299 | 0.163 | 1000 | 1830 | 0.566 | 4.12 |
| 1 | 3.26 | 0.401 | 307 | 2500 | 1.01 | 2.76 |
| 3 | 15.4 | 1.37 | 195 | 2190 | 2.92 | 2.30 |
| 10 | 313 | 16.6 | 32.0 | 602 | 2.21 | 1.03 |
| 50 | 1020 | 601 | 49.2 | 83.1 | 2.16 | 0.900 |
| 200 | 2960 | 4238 | 67.6 | 47.2 | 2.53 | 1.65 |

The pharmacological activity of SZN-043 was evaluated by measuring the mRNA expression levels of Wnt/β-catenin target genes in the liver. RT-qPCR analysis revealed that SZN-043 induced the liver expression of direct Wnt/β-catenin target genes, including *Cyp1a2, Axin2, Notum*, and *Ccnd1*, in a dose-dependent manner (**Figures 3B-3E**). This dose-dependent induction suggests that SZN-043 effectively stimulates Wnt signaling in the liver. Furthermore, because SZN-043 binds to ASGR1 and ASGPR plays a role in serum alkaline phosphatase (ALP) clearance, we investigated whether SZN-043 treatment affected serum ALP levels. We observed that SZN-043 increased serum ALP levels, especially at doses exceeding 10 mg/kg, with a peak at 2-3 days post-dose and a return to baseline by Days 7-9 post-dose (**Figure 3H**). Importantly, there were no dose-dependent differences in liver expression of the tissue-nonspecific ALP gene, *Alpl* **(Figure 3G)**, indicating that the change in serum ALP levels is primarily due to reduced protein turnover resulting from temporary inhibition of ASGR function by SZN-043, rather than alterations in gene expression. Additionally, a dose-dependent increase in expression of the cell proliferation marker gene, *Mki67*, was observed following SZN-043 administration (**Figure 3F**). This finding further supports the proliferative effect of SZN-043 on hepatocytes, suggesting its potential as a therapeutic agent for promoting hepatocyte regeneration in ALD.

### SZN-043 induced a transient wave of proliferation in human hepatocytes

To specifically assess the proliferative capacity of hepatocytes in response to SZN-043, we administered a single dose of SZN-043 at 1, 3 or 10 mg/kg to liver-humanized FRG mice (**Figure S2**) and collected the liver tissues 3 days after dosing. Double immunofluorescence staining with a human-specific ASGR1 (hASGR1) antibody and the proliferation marker Ki67 showed a dose dependent increase in proliferating human hepatocytes when compared to samples exposed to anti-GFP alone. We subsequently dosed SZN-043 daily at 3 and 10 mg/kg to liver-humanized FRG mice for 7 days (**Figure 4A**). Treatment with SZN-043 led to an increase in the number of hASGR1^+^Ki67^+^ double-positive human hepatocytes, as indicated by the yellow nuclei in the merged panel images (**Figure 4B**). This statistically significant increase on Day 2 and Day 3 in double-positive cells was confirmed through quantification using HALO software analysis (**Figure 4C**). Interestingly, prolonged exposure to SZN-043 did not result in a further increase in the percentage of proliferating cells on day 5 and beyond, suggesting that the physiological negative feedback mechanisms preventing uncontrolled proliferation remain intact.

**Figure 4.**
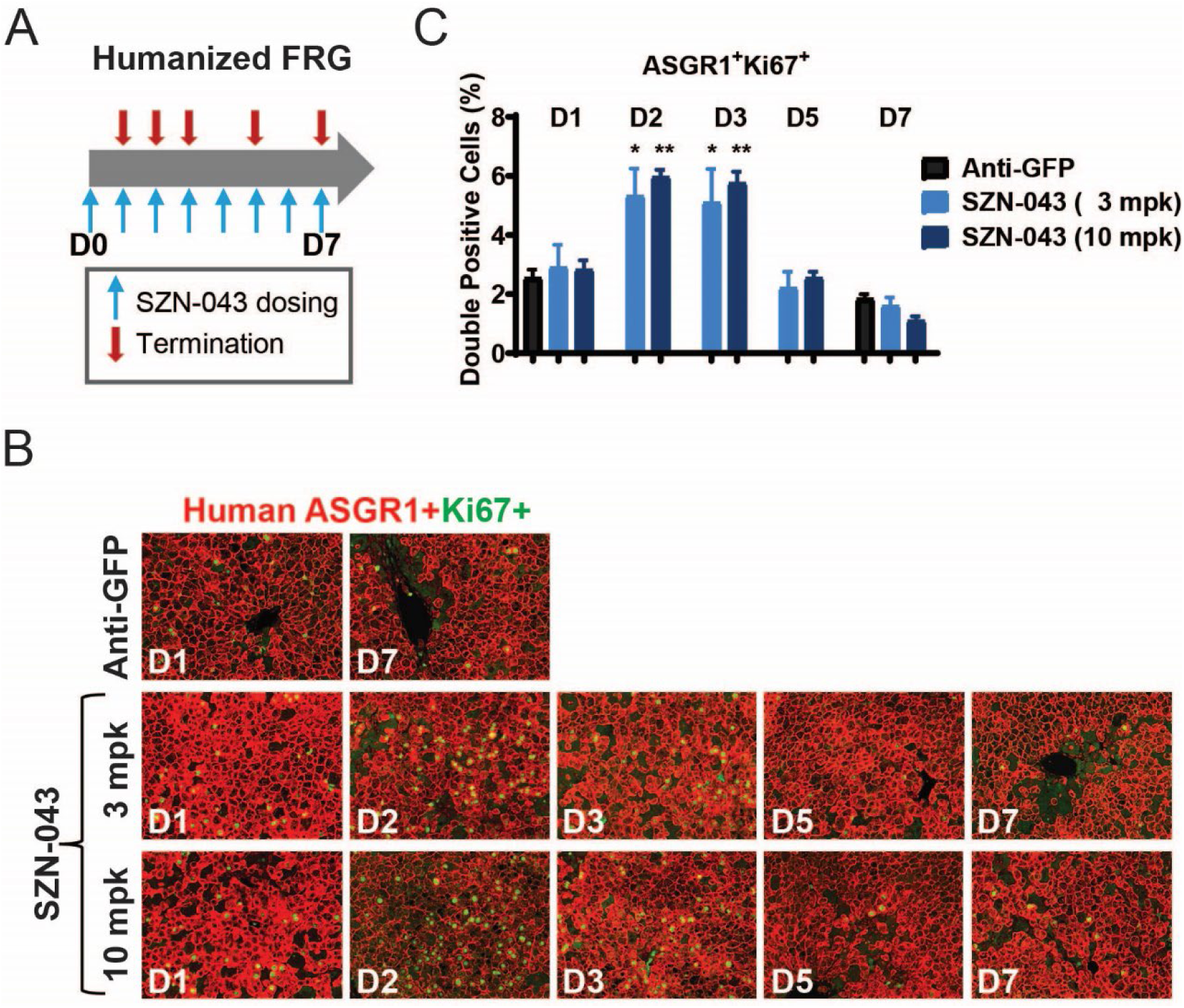
SZN-043 Induced a Transient Wave of Proliferation in Human Hepatocytes. (A) Study design. FRG mice received a single dose, i.p., of either the negative control anti-GFP (10 mg/mL) or the SZN-043 test article at a 3 or 10 mg/kg dose. The left lateral and medial liver lobes were collected upon termination on Day 1, 2, 3, 5 or 7 for immunofluorescence analysis. (B) Left lateral liver tissue sections were doubly stained with the human hepatocyte specific, anti-human ASGR1 (red), and the proliferation marker, anti-Ki67 (green), antibodies. (C) The percentage of doubly positive cells was calculated using HALO imaging analysis software. * p < 0.05, ** p < 0.01

### SZN-043 increased Wnt signaling and transient hepatocyte proliferation in alcohol-induced liver injury models

Accumulating evidence suggest that aged livers are more susceptible to alcohol-induced injury (38). This heightened susceptibility may result from reduced mitochondrial function, age-related alterations in ethanol metabolism, and a diminished capacity for tissue repair (39). To assess the efficacy of SZN-043 in a setting where the spontaneous regenerative capacity of the mouse liver is significantly impaired, we employed a chronic/binge alcohol-induced injury model in aging female B6 mice (**Figure 5**). The mice were fed a 5% ethanol-supplemented liquid diet for 8 weeks and received twice-weekly oral doses of 20% ethanol, except during the first week of the preconditioning period (**Figure 5A**). SZN-043 was administered daily at 30 mg/kg for either three or seven days after the preconditioning period.

**Figure 5.**
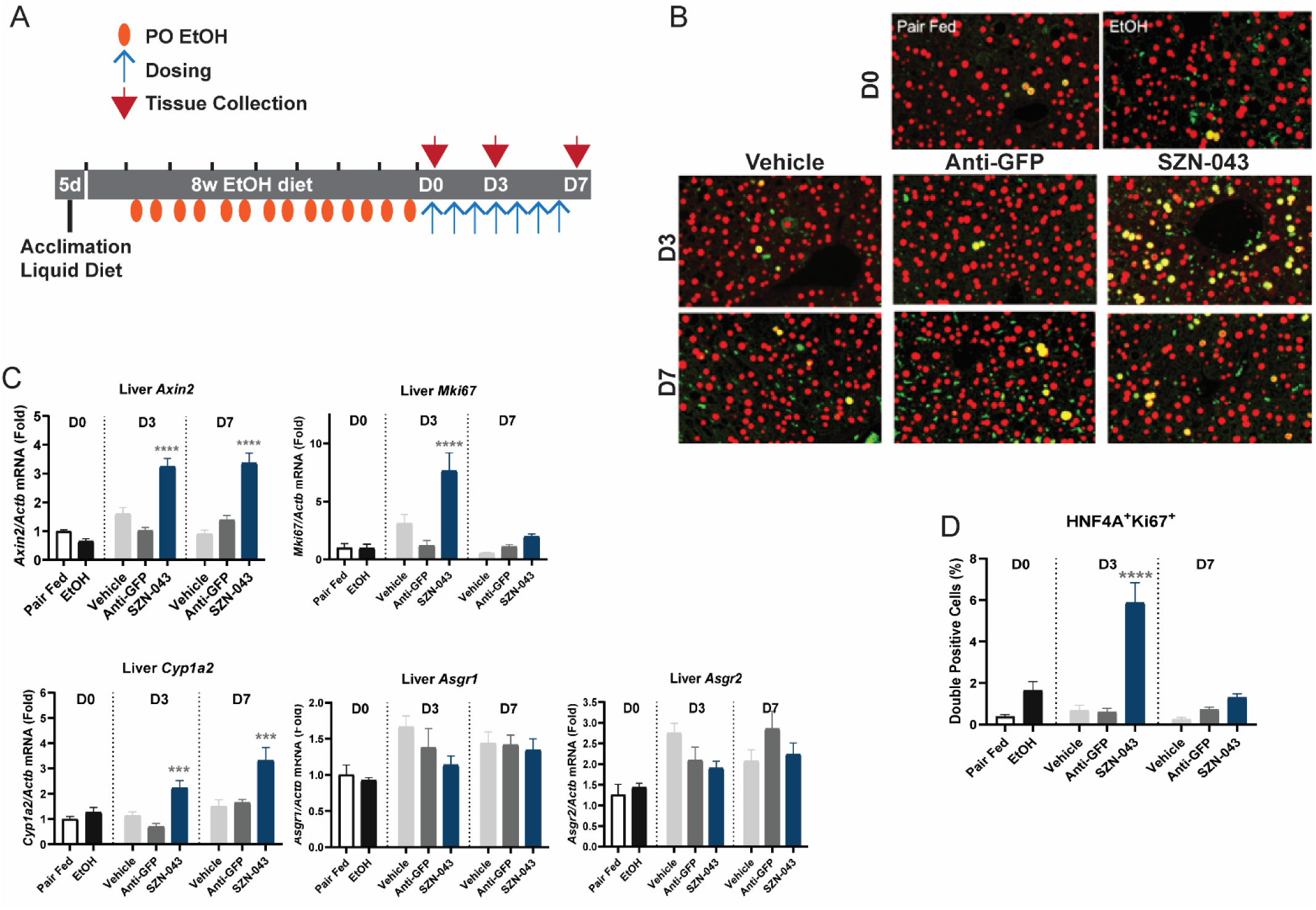
Effect of SZN-043 on Wnt Target Genes and Proliferation Markers in a Chronic-Binge Alcohol-Induced Injury Model in Aging Mice. (A) Study Design. Aging C57BL/6J female mice were acclimated to the Lieber-DeCarli diet for 5 days, followed by an ethanol-supplemented liquid diet for 8 weeks. A control group was pair-fed with an equicaloric diet during this 8-week period. Starting on week 2 of the ethanol-supplemented diet, mice received 20% ethanol by oral gavage twice weekly for the remaining 7 weeks. The pair fed control group received an equivalent volume of ethanol-free equicaloric dose by oral gavage twice weekly also. Ethanol administration was then discontinued, and mice were returned to an ethanol-free liquid diet. The pair-fed control group and an ethanol-fed were euthanized upon ethanol discontinuation, on Day 0, and liver tissue was collected for analysis. Two hours later and daily thereafter until Day 6, mice received either SZN-043 (30 mg/kg), a control antibody anti-GFP (30 mg/kg) or vehicle control. On Day3 and on Day 7, upon termination, liver tissue was collected for analysis. (B) QPCR expression analysis of *Axin2*, *Mki67*, *Cyp1a2*, *Asgr1*and *Asgr2* genes. (C) Immunofluorescence staining with the proliferation marker, anti-Ki67, and the hepatocyte-specific marker, anti-HNF4A antibodies. (D) Percentage of double positive HNF4A and Ki67 cells as determined by quantitative image analysis.

Administration of SZN-043 induced activation of the Wnt signaling pathway, as evidenced by an increase in the expression of Wnt target genes, *Axin2* and *Cyp1a2*, compared to the negative controls (**Figure 5B)**. Additionally, SZN-043 treatment led to the expression of the proliferation marker, *Mki67*, indicating increased hepatocyte proliferation. Immunofluorescence staining confirmed an increase in proliferating hepatocytes, as indicated by double staining for Ki67 and HNF4A, a hepatocyte-specific marker (**Figure 5C and 5D**). Notably, there was minimal spontaneous proliferation in response to ethanol-induced injury compared to that in the pair-fed group. The observed proliferation was transient, with the number of Ki67^+^HNF4A^+^ hepatocytes returning to near baseline levels by day 7 after the start of treatment, despite continued SZN-043 dosing (**Figure 5C and 5D**). Expression levels of *Asgr1* and *Asgr2* were not affected by ethanol or SZN-043 treatment (**Figure 5B**).

In our hands, the use aging mice in this chronic-binge ethanol feeding model failed to reproduce the induction of fibrosis reported in the literature (39). Modest inflammation and steatosis were induced in ethanol-fed mice **(Supplemental Table S3).** SZN-043 did not affect these endpoints significantly. In summary, our findings demonstrated that SZN-043 increased the expression of Wnt target genes and stimulated transient hepatocyte proliferation in a chronic alcohol-induced liver injury aging mouse model.

In a separate study, chronic/binge ethanol treatment in the same model induced significant elevation of *Wnt2* and *Wnt5a* gene expression, along with an upward trend in the expression of *Wnt4* and *Wnt9b* **(Figure S3)**. Conversely, *Wnt6* gene expression was significantly reduced in response to chronic/binge ethanol treatment **(Figure S3B)**. Upon treatment with SZN-043, these trends were reversed. Specifically, *Wnt2* and *Wnt4* were significantly downregulated, and a downwards trend was observed for *Wnt5a* and *Wnt9b* in response to SZN-043 treatment **(Figure S3C)**. These results suggest that SZN-043 may modulate the expression of Wnt signaling pathway components in the context of chronic/binge ethanol-induced liver injury. By reversing the ethanol-induced alterations in Wnt gene expression, SZN-043 may exert its protective effects by restoring the balance of Wnt signaling, which is crucial for liver regeneration and repair processes.

### SZN-043 stimulated hepatocyte proliferation and reduced fibrosis in a CCl_4_-induced liver injury model

Underlying fibrosis is common in patients suffering with alcohol-associated hepatitis and alcohol-associated cirrhosis. To assess whether SZN-043 can induce hepatocyte regeneration in the presence of fibrosis, we conducted experiments in a CCl_4_-induced liver fibrosis model (**Figure 6**). To mitigate the risk of immunogenicity against SZN-043, a human IgG-based protein, we conducted the study in both immunocompetent (B6) and immunodeficient NOD. CB17-Prkdc*^scid^*/J (SCID) mice. Mice were subjected to CCl4 treatment to induce liver fibrosis over 10 weeks. Interestingly, both B6 and SCID mouse strains exhibited similar levels of fibrosis after this treatment (**Figure 6B and 6D**). Following this induction phase, the mice were treated with either SZN-043, RSPO, or an isotype antibody control (anti-βGal) for 14 days. Treatment with SZN-043 led to an increase in the expression of the liver proliferation marker *Mki67* after 14 days of treatment in both mouse strains, indicating enhanced hepatocyte proliferation (**Figure 6A**). Immunofluorescence staining revealed that SZN-043 specifically induced Ki67 expression in HNF4A+ hepatocytes in both strains, suggesting hepatocyte expansion in response to liver injury induced by CCl_4_ (**Figure 6B)**. Moreover, the regenerative efficacy of SZN-043 was comparable to that of RSPO (**Figure 6A and 6C**). Additionally, treatment with SZN-043 resulted in a modest but significant reduction in the fibrotic area in the liver compared to the control group, as demonstrated by picrosirius red (PSR) staining and quantification of liver sections (**Figure 6B and 6D**). Overall, these findings indicate that SZN-043 stimulates hepatocyte-specific cell regeneration even in the presence of fibrosis induced by CCl_4_ in both strains of mice. Furthermore, this regeneration was associated with a modest but significant improvement in liver fibrosis, suggesting that Wnt activation targeted to hepatocytes does not stimulate fibrogenesis.

**Figure 6.**
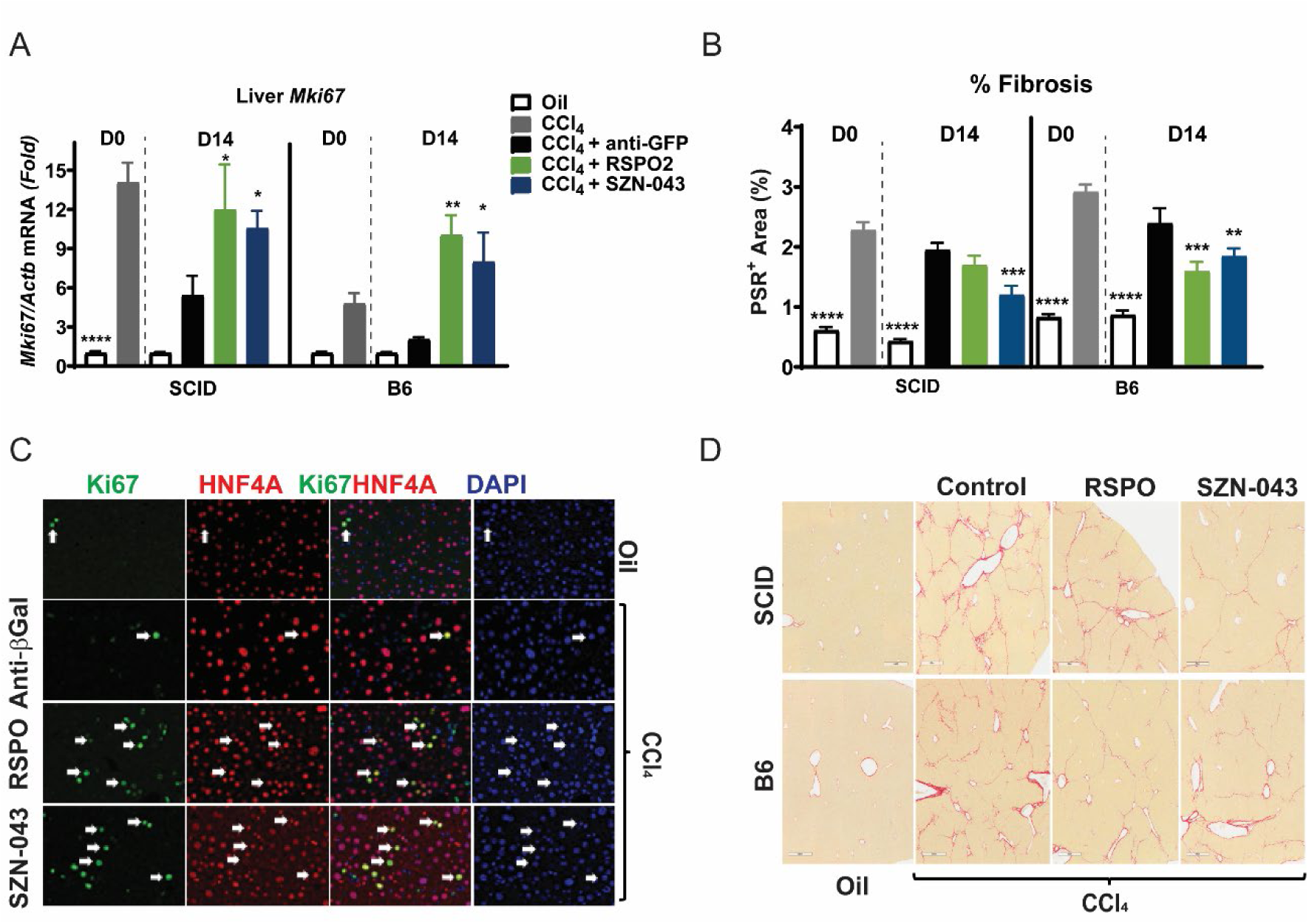

## DISCUSSION

ALD encompasses a range of liver conditions, including steatohepatitis, fibrosis, and cirrhosis. Severe AH, often characterized by a MELD score greater than 20, is frequently associated with underlying cirrhosis (3). These cases are marked by impaired hepatocyte proliferation and have a significant mortality rate, with approximately 15 to 20% mortality at one month and 30% mortality at three months (40). The deficiency in hepatocyte proliferation observed in AH provides an opportunity for intervention with a therapeutic agent capable of inducing a regenerative process to enhance hepatic function and improve patient outcomes (41). However, there is a notable gap in the approved pharmacotherapeutics for the treatment of sAH. Current treatment options are limited, with corticosteroids being the only class of drugs recommended for sAH. However, their efficacy is modest, resulting in only slight reductions in mortality, and only a subset of patients is eligible for this treatment (3, 42).

The Wnt signaling pathway, along with its regulator RSPO, plays crucial roles in liver regeneration (33). In this study, we investigated the expression of key Wnt pathway-related genes in two cohorts of patients with ALD. Our findings demonstrated upregulation of *WNT2B, WNT4, WNT7B, WNT9A, WNT10A, and WNT10B* in cirrhotic livers, whereas *WNT2* and *WNT11* were downregulated. Interestingly, a recent study by Kim et al. reported similar changes in the expression of these Wnt ligands in patients with sAH, except for *WNT11 and FZD6*, which was upregulated in patient samples (43). Consistent with these findings, we also observed the upregulation of *FZD6* in our cohort of patients with alcohol-associated cirrhosis, although the difference was not statistically significant. Collectively, these results suggest that sAH and alcohol-associated cirrhosis share extensive features of WNT ligand dysregulation.

The role of individual WNT family members in liver homeostasis is not fully understood, but there is evidence suggesting that WNT4 acts as an antagonist through a non-canonical pathway, while other Wnt family members identified in our study act as agonists through canonical pathway. The findings from our study suggest a reduction in Wnt activity in patients with ALD, as indicated by the lack of significant elevation in proliferation marker genes in the alcohol-associated cirrhosis cohorts. Interestingly, the expression of *RSPO3*, the main R-spondin in the liver, and E3 ligases remained unchanged in ALD patients. However, there was a notable decrease in the expression of *LGR4* and *LGR5*, which are crucial co-receptors for R-spondin-mediated Wnt signaling in the liver. The downregulation of LGR receptors could potentially impair the regenerative response mediated by R-spondins and other Wnt-mediated functions in ALD. Nevertheless, it is important to note that the activity of SZN-043, a hepatocyte-targeted Wnt mimetic, is not dependent on the presence of LGR co-receptors. Instead, SZN-043 specifically binds to hepatocytes via ASGR1 and engages the E3 ligases in an LGR-independent manner (28). Therefore, the reduction in *LGR4* expression in patients with ALD should not impede the activity of SZN-043. Moreover, the preservation of ASGR1 expression in these patients should facilitate the targeting of SZN-043 to hepatocytes, potentially restoring Wnt signaling and promoting hepatocyte regeneration in ALD. This highlights the potential of SZN-043 as a therapeutic agent for ALD, despite the observed dysregulation of LGR receptors in these patients.

This dichotomy in the effects of WNT family members, with some acting as agonists and others as antagonists, makes it challenging to predict the combined impact of their dysregulation. Further research is needed to clarify the specific roles of each family member on different hepatic cell types. Despite the complexity of WNT signaling, our study revealed a significant downregulation of several hepatocyte-specific or -enriched Wnt target genes, *CYP1A2, CYP2E1, OAT, GLUL, LGR5, SZNRF3*, in liver samples from patients with ALD, as evidenced by multiple analyses including immunostaining, qPCR, and bulk RNAseq. This suggests that, overall, there is a loss rather than an increase in Wnt activity in ALD liver samples. Some Wnt target genes expressed in epithelial and mesenchymal liver cell populations, such as *AXIN2*, *CCND1* and *NOTUM* were not consistently and significantly changed. This may be due to the large increase in mesenchymal cells in ALD tissue samples confounding the results. No significant increase in the expression of these genes were observed either, consistent with the lack of Wnt activity increase in these samples. R-spondins have been reported to act via additional pathways in addition to the Wnt signaling pathway (for review (44)). Here, we cannot exclude the R-spondin mimetic, SZN-043 binds to additional receptors than ZNRF3 and RNF43 in hepatocytes. It remains to be tested whether the mutations introduced in SZN-043 to prevent its binding to LGR5, also affect its binding to other potential co-receptors such as FGFR4.

The pharmacological properties of SZN-043 were extensively characterized using various in vivo models, including both non-injury models in mouse and human hepatocytes, as well as ethanol-induced injury and fibrosis mouse models. One crucial aspect assessed was the clearance and mean residence time of SZN-043, which decreased with increasing dose. This observation is consistent with SZN-043’s binding to ASGR1, a highly abundant protein enriched in hepatocytes, and subsequent target-mediated drug degradation. This finding was further supported by the concurrent increase in serum ALP levels, with little or no corresponding change in the expression of *Alpl*, a gene encoding alkaline phosphatase. Elevated serum ALP levels are indicative of the removal of ASGR1 from the hepatocyte surface by SZN-043, as ASGR1 plays a critical role in ALP clearance (45). Importantly, a similar ALP elevation is also observed in human carriers of *ASGR1* loss-of-function variants and *ASGR1* knockout mice (45, 46). Furthermore, the increase in serum ALP levels may serve as a useful non-invasive biomarker of ASGR1 target engagement in future nonclinical and clinical studies involving SZN-043. This comprehensive characterization of SZN-043’s pharmacological properties provides valuable insights into its mechanism of action and safety profile, supporting its potential as a therapeutic agent for ALD.

A substantial number of publications report an association of ASGR1 with potential adverse events in animal models (for review (47)), although the conclusions of these datasets are not always translated across species and even sometimes inconsistent within the same species (48, 49). In this study, we focused on the SZN-043 efficacy and potential mode of action in preclinical models. As part of a clinical development program, separate safety assessment studies are ongoing in relevant species following standard approaches.

Activation of Wnt signaling by SZN-043 in the liver exhibited a dose-dependent response, with induction occurring even at low doses of 0.3 mg/kg. Interestingly, there was a dose-dependent delay in reaching the peak induction of several Wnt target genes, suggesting a right-shift in the dose-response curve. For instance, while lower doses of SZN-043 induced maximal *Axin2* expression at 24 hours post-dosing, higher doses (200 mg/kg) showed peak efficacy at 7 days post-dosing. This delay in peak efficacy could potentially result from the rapid depletion of ASGR1 on the hepatocyte cell surface, followed by the time needed for ASGR1 recycling.

We previously reported that the induction of Wnt signaling by SZN-043 was accompanied by a substantial increase in hepatocyte proliferation in healthy and thioacetamide-induced fibrotic mice (28). Here we report that SZN-043 induces hepatocytes proliferation in humanized livers, in an aging alcohol-induced injury mouse model, and in a CCl_4_-induced fibrosis mouse model. Importantly, this proliferation was transient, indicating the presence of physiological Wnt-induced negative feedback loop mechanisms. Notably, the expression of *Axin2* and *Notum*, key negative regulators of Wnt signaling, remained elevated even in the presence of increased proliferation. AXIN2 promotes the degradation of β-catenin, thereby inhibiting Wnt signaling, whereas NOTUM acts as an extracellular de-acylase, inactivating Wnt ligands. The maintenance of elevated *Axin2* and *Notum* expression suggests that treatment with SZN-043 does not lead to uncontrolled proliferation, as the negative regulatory mechanisms of Wnt signaling remain intact. Overall, these findings provide important insights into the regulatory dynamics of Wnt signaling induced by SZN-043 and their implications for hepatocyte proliferation.

The efficacy of SZN-043 in inducing proliferation of human hepatocytes in FRG liver humanized mice at doses of 3 and 10 mg/kg, which was also effective in inducing proliferation in mouse hepatocytes, underscores its potential therapeutic utility. While only hepatocytes in FRG mice are of human origin, the model may not fully recapitulate the physiological conditions of a human liver due to the continuous cell death of mouse hepatocytes caused by the Fah-KO mutation. This ongoing cell death could potentially impact the liver environment and influence the interpretation of results. Nevertheless, these findings demonstrate that human hepatocytes possess the machinery necessary to respond to SZN-043 activity, suggesting translational relevance. Moving forward, the development of more complex humanized liver models incorporating both parenchymal and non-parenchymal hepatic cells of human origin, such as the recently developed model (50), will likely provide valuable insights into the mechanisms underlying the action of SZN-043 in human livers. These advanced models hold promise for elucidating the interplay between different cell types within the liver microenvironment and their responses to Wnt pathway modulation by SZN-043, thereby facilitating further preclinical investigations and potentially informing clinical translation.

The efficacy of SZN-043 in stimulating Wnt activation and hepatocyte proliferation in a chronic-binge aging mouse model of alcohol-associated liver injury highlights its potential therapeutic utility in situations where hepatic regenerative capacity is compromised. Notably, this study utilized older female mice, which exhibit reduced regenerative capacity and heightened susceptibility to alcohol-induced liver injury, reflecting aspects of human pathology (39). Interestingly, alcohol-induced liver injury did not affect the expression of Asgr1, the target receptor for SZN-043, suggesting that the mechanism of action of SZN-043 remains viable in the context of AH. Furthermore, SZN-043 administration resulted in the upregulation of Wnt-target genes, such as *Cyp1a2* and *Axin2*, indicative of enhanced Wnt pathway activity, along with the induction of proliferation markers such as cyclin D1 and *Mki67* (Ki67). It is important to note that the observed proliferation was transient, returning to baseline levels by day 7, whereas the elevation in Wnt target gene expression persisted at this later time point. This temporal discrepancy suggests that while SZN-043 treatment induces a finite wave of hepatocyte proliferation, the negative feedback mechanisms inherent in Wnt signaling effectively regulate the duration of this proliferative response, ensuring controlled and physiological regeneration.

A limitation to these studies is the modest amount of liver-injury observed in mouse models when compared to those observed in ALD patients. To date, there is no model that reproduce consistently the severe damage observed in alcohol-associated cirrhosis along with elevated MELD score. The combination of the chronic-binge ethanol treatment with a high-fat, high-carbohydrate diet, may have better mirrored the metabolic challenges seen in human ALD (51, 52). A combination of acetaminophen, LPS and CCl_4_ treatments was suggested to model end-stage multi-organ liver disease (53). In our hands, these treatments failed to induce liver injuries greater than with CCl_4_ alone (unpublished results). Alternatively, combining ethanol with CCl4 could also lead to more severe fibrotic livers. All of these models result in similar pericentral fibrosis with central-to-central bridging.

The CCl_4_-induced liver fibrosis model employed here remains a commonly used and robust murine model of fibrosis, despite not reaching the severity of fibrosis observed in humans. Treatment with SZN-043 in the CCl_4_-induced fibrosis model activated Wnt signaling and stimulated hepatocyte proliferation, while concurrently reducing the fibrotic area, as indicated by (PSR) staining. The effect of SZN-043 on fibrosis was not further improved in immunodeficient mice when compared to B6 mice, arguing that the modest but significant effect size was not resulting from neutralizing anti-drug antibodies. The exact mechanism underlying this improvement in fibrosis remains unclear and may involve intricate interactions within the liver microenvironment. For example, Wnt activation is known to induce the secretion of hepatokines, such as LECT2, which in turn can influence macrophage activity (54, 55) While some studies have suggested that Wnt activation may exacerbate fibrosis by stimulating stellate cells (56), others reported a protective role of RSPO in preventing fibrosis (24). The observed reduction in fibrosis following SZN-043 treatment may be attributed to hepatocyte proliferation induced by SZN-043, leading to the transformation of the liver microenvironment from a pro-inflammatory and profibrotic state to an anti-inflammatory and anti-fibrotic state (57, 58). This hypothesis is consistent with the concept of hepatocyte-driven regulation of the liver microenvironment, wherein changes in hepatocyte behavior can impact the activation and function of other liver cell populations, including stellate cells. However, the complex interplay between different liver cell populations and the specific roles of individual Wnt and Fzd family members in fibrogenesis warrants further investigation.

An anti-RSPO3 antibody, rosmantuzumab (OMP-131R10), was previously developed and tested for the treatment of metastatic colorectal cancer(59, 60). In contrast to SZN-043, OMP-131R10 was developed to inhibit rather than enhance RSPO activity. In a phase 1 clinical trial, high doses of OMP-131R10 reduced the osteogenesis marker P1NP, showing the multi-organ effects that non-specific RSPO-based therapies may entail.

By selectively targeting hepatocytes and activating the Wnt signaling pathway, SZN-043 constitute a potential therapeutic candidate for treating sAH. Here we demonstrated robust pharmacological effects, promoting hepatocyte proliferation and potentially attenuating liver fibrosis. Given the limitations associated with the current treatment options for sAH, such as limited efficacy and eligibility criteria, there is a clear unmet need for novel therapeutic approaches. SZN-043 has the potential to address this need by targeting the critical molecular pathways involved in liver regeneration and repair. Furthermore, the specific mechanism of action of SZN-043, which circumvents the requirement for LGR co-receptors and instead targets the highly abundant ASGR1 receptor on hepatocytes, offers advantages in terms of specificity and efficacy. This targeted approach minimized potential off-target effects and enhanced the therapeutic potential of SZN-043 in such patients. Overall, the results presented here support further investigation of SZN-043 as a potential treatment option for sAH. Future studies, including clinical trials, will be crucial to evaluate the safety and efficacy of SZN-043 in patients and assess its potential impact on improving outcomes in this challenging disease. Continued research efforts aimed at elucidating the underlying mechanisms of SZN-043’s therapeutic effects and optimizing its clinical application hold promise for advancing the management of sAH and improving patient outcomes.

## Author contributions

Helene Baribault, Trevor Fisher, Mehaben Patel and Geertrui F. Vanhove conceptualized and supervised the studies. Helene Baribault drafted and finalized the manuscript, Trevor Fisher, Mehaben Patel, Shalaka Deshmukh, Darshini Shah, Chenggang Lu, Maureen Newman, Jay Ye and Zhiyong Yang performed experiments, Geertrui F. Vanhove, Russell Fletcher, Yang Li and Jay Tibbitts contributed to the discussion and interpretation of results and reviewed the manuscript. Suthat Liangpunsakul provided comments, reviewed, and edited the manuscript.

## Ethics statement

Human liver tissue remnants from liver resections and explants were used for histological processing and total mRNA extraction. The ethics committee of the Hospital Clinic de Barcelona approved the study protocol (HCB/2018/0028), and the Institutional Review Board at Indiana University. In all cases, patients signed an informed consent before resection or transplantation.

## ABBREVIATIONS

ALD: Alcohol-associated liver disease
AH: Alcohol-associated hepatitis
B6: C57BL/6J
CCl_4_: Carbon tetrachloride
EtOH: Ethanol
FDR: False discovery rate
FDRsig: Statistically significant FDR change
FRG: FRG® KO (mouse strain lacking FAH, RAG2 and IL2RG)
GFP: Green fluorescent protein
HSC: Hepatic stellate cell
L-D: Lieber-DeCarli
MELD: Model for end-stage liver disease
n.a.: Not applicable
PSR: Picrosirius Red
RSPO: R-spondin
sAH: Severe alcohol-associated hepatitis
SCID: NOD.CB17-Prkdc*^scid^*/J
SEM: Standard error of the mean

## ACKNOWLEDGMENTS

This work was supported by Surrozen Inc., South San Francisco, CA, USA. We are grateful to Jordi Garcia from Barcelona Liver Biosciences for services and discussions. We thank Dr. Vincent Meador (Pacific Tox Pathology) for his continued support in pathological analysis. We are grateful to Kristofor Feger and Tiep Le for their assistance with animal care.

## SUPPLEMENTARY MATERIAL

### EXPERIMENTAL PROCEDURES

#### Transcriptome Analysis of Livers from Alcoholic Cirrhosis Patients

Human liver tissue remnants from liver resections and explants from patients with alcohol-associated liver disease were used for histological processing and total mRNA extraction. Non-tumorous liver tissue from hepatic resections was used as a healthy control and compared to liver explants from patients with alcohol-associated liver disease (n=12 per group), (**Supplementary Table S1**). The following genes encoding Wnt and RSPO ligands, FZD receptors, RSPO and SZN-043 target receptors, and Wnt target genes were analyzed by qPCR using the following probes from Applied Biosystems: *ASGR1* (Hs01005019_m1), *ASGR2* (Hs00154160_m1), *CYP1A2* (Hs00167927_m1), *FZD1* (Hs00268943_s1), *FZD2* (Hs00361432_s1), *FZD3* (Hs00907280_m1), *FZD4* (Hs00201853_m1), *FZD5* (Hs00361869_g1), *FZD6* (Hs01095627_m1), *FZD7* (Hs00275833_s1), *FZD8* (Hs00259040_s1), *FZD9* (Hs00268954_s1), *FZD10* (Hs04999826_s1), *RNF43* (Hs00214886_m1), *RSPO1* (Hs00543475_m1), *RSPO2* (Hs04400416_m1), *RSPO3* (Hs00262176_m1), *RSPO4* (Hs01382765_m1), *WNT1* (Hs00180529_m1), *WNT2* (Hs00608224_m1), *WNT2B* (Hs00921614_m1), *WNT3* (Hs00902257_m1), *WNT4* (Hs01573505_m1), *WNT5A* (Hs00998537_m1), *WNT5B* (Hs01086864_m1), *WNT7A* (Hs01114990_m1), *WNT7B* (Hs00536497_m1), *WNT8B* (Hs00610126_m1), *WNT9A* (Hs01573829_m1), *WNT9B* (Hs00287409_m1), *WNT11* (Hs01045906_m1), and *ZNRF3* (Hs00393094_m1).

Additionally, global transcriptome profiling of human liver tissue was performed using RNA-Seq. Sequencing libraries were prepared following the TruSeq Stranded mRNA Sample Preparation Guide, employing the TruSeq Stranded mRNA Library Prep kit and TruSeq RNA Single Indexes (Illumina Inc.). The libraries were subsequently sequenced in a HiSeq2500 System (Illumina) with a single-end type read and a read length of 101 nucleotides, generating 36 million reads. This comprehensive approach allowed for the examination of gene expression profiles and identification of potential dysregulations in the Wnt/β-catenin pathways in alcohol-associated liver disease. The expression levels of SZN-043 target receptors and other molecular components of the Wnt/β-catenin pathways were also performed and validated in the liver tissues of another cohort of healthy controls (N=5) and patients with alcohol-associated cirrhosis (N=4) **(Supplementary Table S2)**.

#### Histological analysis of the liver tissues

Paraffin-embedded liver sections were utilized for histological analyses, including Picrosirius Red staining and immunohistochemical detection of specific proteins. For immunohistochemical detection of CYP1A and ASGR1 specific antibodies were used. A rabbit anti-CYP1A2 antibody from BosterBio (Catalog # PB9545) and a rabbit anti-ASGR1 antibody from ThermoFisher Scientific (Catalog # PA5-80356) were employed. After incubation with the primary antibodies, the sections were incubated with a biotinylated secondary antibody, such as a biotinylated goat anti-rabbit IgG antibody from Abcam (Catalog # ab6720).

#### Recombinant proteins

SZN-043 is a modified version of αASGR1-RSPO2-RA (28), which is a humanized IgG1 monoclonal antibody (mAb) with complementarity-determining regions (CDRs) against ASGR1, fused to a mutated RSPO2 on each N-terminus of the heavy chain (28). To eliminate crystallizable fragment (Fc) effector function, the Fc region of SZN-043 has been engineered with an N297G mutation (61). Additionally, to prevent LGR binding, point mutations (F105R and F109A) were introduced within the Fu2 domain of RSPO2 (28). SZN-043 differs from αASGR1-RSPO2-RA by 6 amino acids, including at the positions F105E and F109E. These changes ensure that SZN-043 retains its binding and pharmacological effects while optimizing its drug-like properties properties, such as specificity and stability. An additional variant of SZN-043, SZN-043.v2, was developed, which includes an S51A replacement. Both variants of SZN-043 (SZN-043 and SZN-043.v2) exhibit similar binding affinity for human ASR1 (with dissociation constants K_D_ of 4.84E-8 and 6.83E-8 M, respectively) and comparable potency as determined by SuperTopFlash luciferase reporter assay in the human liver cell line Huh-7. RSPO refers to the fusion Fc-RSPO2 protein described previously (28), where RSPO2 (S36-E143) is fused to the C-terminus of human IgG1. Similarly, the anti-GFP protein mentioned was also described in (28).

#### Animals

The animal experimentation conducted in this study adhered to the criteria outlined in the Guide for the Care and Use of Laboratory Animals, 8th edition, prepared by the National Academy of Sciences. Protocols for animal experimentation were approved by the Surrozen Operating, Inc. Institutional Animal Care and Use Committee. Upon arrival, mice were housed in compliance with animal welfare standards using the Innovive Disposable IVC Rodent Caging System for mice (Innovive, San Diego, CA). They were acclimatized for at least two days before the commencement of the study. During the study period, mice had ad libitum access to food (2018 Teklad global 18% protein rodent diet, Envigo, Indianapolis, IN) and purified, laboratory-grade acidified water. Termination procedures were conducted in accordance with the current requirements of the Guide for the Care and Use of Laboratory Animals, 8th edition, and the American Veterinary Medical Association (AVMA) Guidelines on Euthanasia. At termination, mice were anesthetized using isoflurane, and blood was collected via cardiac puncture for serum and plasma collection. Subsequently, euthanasia was performed via cervical dislocation. Liver samples were collected post-euthanasia and subjected to different analyses. Some liver samples were formalin-fixed and paraffin-embedded for immunofluorescence and histopathological analysis. Additionally, other liver samples were snap-frozen in liquid nitrogen and stored at -80°C for subsequent gene expression analyses.

#### Dose Response Study

In this study, 8-week-old C57BL/6J male mice obtained from Jackson Laboratories were utilized. The mice were divided into a control group (n=6) and groups receiving different doses of SZN-043 (0.3, 1, 3, 10, 50, or 200 mg/kg), with 18 mice per dose group (n=18/group). Mice in the control group did not receive any treatment, while those in the treatment groups (totaling 108 mice) received a single intravenous (IV) bolus dose of SZN-043 according to their assigned treatment group. All mice were monitored for up to 14 days following dosing. Blood samples were collected from a subset of mice (n=3 from each dose group) at specified time points (5 minutes, 15 minutes, 45 minutes, 2 hours, 6 hours, 24 hours, 2 days, 3 days, 4 days, 7 days, 9 days, and 14 days after dosing) for quantitation of serum SZN-043 concentrations and biomarker analyses. Each mouse was bled a maximum of twice to adhere to recommended blood collection volume guidelines(62). At the end of the study period, livers were collected from all mice for gene expression analysis, providing insights into the effects of SZN-043 treatment on hepatic gene expression profiles. This experimental design allowed for the comprehensive evaluation of SZN-043 pharmacokinetics, biomarker responses, and potential effects on hepatic gene expression over a 14-day period across various dosages in a preclinical mouse model.

#### Human hepatocyte proliferation in FRG mice

In this study, FRG mice, which are immunocompromised due to genetic deficiencies in *Rag2* and *Il2rg*, were utilized. These mice lack the tyrosine catabolic enzyme fumarylacetoacetate hydrolase encoded by the *Fah* gene, resulting in the accumulation of toxic levels of fumarylacetoacetate and maleylacetoacetate in endogenous mouse hepatocytes. To control the build-up of these toxic catabolites, the mice were maintained on a low tyrosine diet and treated with the tyrosine metabolism inhibitor NTBC throughout their lives. The FRG mice used in this study were male, 22 to 25 weeks old, on a C57BL/6 background, and were repopulated with human hepatocytes (70%+) obtained from Yecuris (Tualatin, OR, Catalog # 10-0006). Upon arrival, the NTBC-free FRG mice were housed in the clean room of the vivarium due to their severe immunodeficiency. They were acclimated for a minimum of two days prior to the start of the study and housed using the Innovive Disposable IVC Rodent Caging System for mice (Innovive, San Diego, CA). The mice had ad libitum access to a low tyrosine (0.55%) and high-fat-content (11%) diet (PicoLab® High Energy Irradiated Diet 5LJ5, Newco, Hayward, CA) and purified, laboratory-grade acidified water. The low tyrosine diet was essential to prevent the build-up of toxic metabolites in the FRG mice due to their Fah deficiency.

The mice were randomized into treatment groups based on body weight and serum human albumin concentrations provided by the vendor. All groups received a single dose of either SZN-043 (3 mg/kg or 10 mg/kg) or a negative control anti-GFP 10 mg/kg via intraperitoneal injection on Day 0 (0 hour). Mice were euthanized for tissue collection at specified time points after the start of treatment (Day 2, 3, 5, or 7). The extent of human hepatocyte contribution to livers was confirmed using anti-human specific ASGR1 immunostaining of histological sections.

#### Alcohol-induced liver injury mouse model

In this study, the mouse model of chronic-binge ethanol feeding (the NIAAA model) (63), was used with 13.5 months old aging female C57BL/6J mice (Jackson Laboratories, Bar Harbor, MA). The mice, group-housed with up to four mice per cage, were transferred to rat cages from Innovive (#RVX1) and were fed either a 5% ethanol or equicaloric control diet for the duration of the study, as previously described. Liquid diets were provided using two 50 mL liquid diet feeding tubes and holders from Bio-Serv (#9019, #9015) per cage, with feeding tubes changed daily. Binges of ethanol or equicaloric maltose dextrin were administered to mice by oral gavage, twice weekly. The ethanol solution (20% v/v) was prepared immediately before use by mixing 4.2 mL of 95% ethanol with 15.8 mL water and administered to ethanol-treated mice at 5 mL/kg. The maltose dextrin solution (28.6% w/v) was prepared immediately before use by mixing 5.9 g of maltose with water for a final volume of 20 mL and administered to pair-fed mice at 5 mL/kg. Daily food consumption per cage was recorded to calculate the average food consumption per mouse during the ethanol-feeding period. Body weights were collected and recorded twice weekly throughout the study, with the final weight recorded at termination.

In this particular study, 14-month-old mice in the ethanol-fed groups were given free access to the Lieber-DeCarli (L-D) diet containing 5% (v/v) ethanol for eight weeks, while the pair-fed group received the L-D control diet with maltose dextrin substituting for ethanol. Starting in the second week and continuing through week 8 of the feeding period, ethanol-fed mice received twice-weekly gavage doses of 20% ethanol for a total of 14 gavages. After the ethanol feeding period, mice were returned to the L-D control diet and randomized into treatment groups (n = 6 to 8 mice/group). The pair-fed and ethanol-fed groups were terminated on Day 0. Four hours after discontinuation of ethanol feeding, the remaining mice were injected intraperitoneally either daily with SZN-043 (30 mg/kg) or twice weekly with negative controls vehicle and anti-green fluorescent protein (GFP) (30 mg/kg) groups and terminated at Day 3 or Day 7 after the start of dosing. Liver samples were collected for analysis.

#### Liver fibrosis mouse model

In the CCl_4_-induced liver injury model, 32 SCID and 32 B6 male mice obtained from Jackson Laboratories were utilized. These mice were injected intraperitoneally (IP) with CCl_4_ (0.5 mL/kg) diluted in an olive oil carrier (CCl_4_:olive oil; 1:3 ratio v/v) twice weekly for a total of 10 weeks to induce hepatic fibrosis. A control group received IP injections of only the olive oil carrier (0.5 mL/kg), also administered twice weekly for 10 weeks. Two days after discontinuation of CCl_4_ administration, mice from both strains were randomized, based on body weight, into one of three treatment groups (n = 8/group). The treatment groups included: (i) SZN-034.v2 at a dose of 20 mg/kg once daily (active treatment group), (ii) RSP02 at a dose of 4.6 mg/kg twice weekly (positive control), and anti-βGal antibody at a dose of 10 mg/kg twice weekly (negative control). Each test article was administered IP as scheduled over a period of 2 weeks. Additionally, the remaining 8 mice that received only olive oil in the liver injury phase served as the vehicle control group, and no additional treatments were administered to this group. Upon termination, liver samples were collected from all mice for analysis.

#### Serum Chemistry

Blood samples were collected in serum separation tubes with gel from Thermo Fisher Scientific (Waltham, MA, 22-030-401). After collection, the tubes were centrifuged at 10,000 RPM for 7 minutes. Following centrifugation, the supernatant (serum) was carefully transferred to new tubes. These serum samples were then stored at -20°C until analysis. Serum alkaline phosphatase (ALP) levels were analyzed using a Vet Axcel® chemistry analyzer from Alfa-Wasserman Diagnostic Technologies, LLC (West Caufield, NJ #404200-3), along with alkaline phosphatase test reagents (SA2002).

#### Enzyme-linked immunosorbent assay (ELISA)

The analysis of serum samples for concentrations of SZN-043 was performed using an anti-idiotype assay in conjunction with the MSD GOLD Streptavidin SECTOR Plate (MSD, #L15SA-1). In this assay, a biotinylated neutralizing anti-idiotype antibody (designated as 8#13) was chosen to capture the ASGR1-binding regions of SZN-043. Additionally, a ruthenylated-labeled neutralizing anti-RSPO2 antibody (designated as 3A3D7) was selected to detect the E3 ligase-binding regions of SZN-043. To initiate the assay, tripropylamine (TPA, MSD Gold Read Buffer) was added to the plate. Upon the application of an electric charge, an electrochemiluminescent signal was generated and detected by the MSD instrument. This signal is proportional to the concentration of SZN-043 present in the serum sample. The back-calculated concentrations (BCC) of the analyte were determined using a 4-parameter logistic (4PL) fit model with 1/Y2 weighting. This analytical approach allows for the accurate quantification of SZN-043 concentrations in serum samples. The bioanalytical assay strategy for measuring SZN-043 concentrations in serum was developed in accordance with M10 Bioanalytical Method Validation and FDA Bioanalytical Method Validation Guidance for Industry (2018) guidelines.

#### Histological and Immunofluorescence Analysis

The livers obtained from each mouse underwent fixation with 10% neutral buffered formalin for 24 hours, after which they were transferred to 70% ethanol for preservation. Subsequently, the livers were trimmed, processed into paraffin blocks, and sectioned to prepare histological slides. For histological analysis, the sections were stained with PicroSirius Red (PSR), a dye commonly used to visualize collagen deposits and fibrosis. The stained sections were then sent to a board-certified veterinary pathologist, Vincent Meador, DVM, ACVP, at Pacific Tox Path, LLC, for blinded evaluation. Blinded evaluation ensures unbiased assessment of the histological features by the pathologist. To quantify the fibrotic area, Image J software from the National Institutes of Health (Bethesda, MD) was utilized. This software allows for the quantification of the percentage of tissue stained positive for PSR on whole liver sections, providing an objective measure of fibrosis severity. For double immunofluorescence staining of Ki67 and HNF4A, primary antibodies including rat anti-Ki67 antibodies (Thermo Fisher Scientific, 50245564) and rabbit monoclonal anti-HNF4A-N-terminal antibodies (Abcam, ab199431) were used. Subsequently, secondary antibodies including goat anti-rat IgG H&L secondary antibody (Abcam, Alexa Fluor® 488, ab150157) and donkey anti-rabbit IgG H&L secondary antibody (Abcam, Alexa Fluor® 647, ab150075) were applied. The labeled sections were then imaged using a Leica DMi8 microscope equipped with a DFC7000T camera, allowing for visualization and analysis of Ki67 and HNF4A expression within the liver tissue.

For the double immunofluorescence staining of Ki67 and human ASGR1, the following antibodies were utilized: rat anti-Ki67 antibodies from Thermo Fisher Scientific (50245564) and rabbit anti-human ASGR1 antibodies from Thermo Fisher Scientific (PA5-80356). After primary antibody incubation, secondary antibodies including goat anti-rat IgG H&L secondary antibody (Abcam, Alexa Fluor® 488, ab150157) and donkey anti-rabbit IgG H&L secondary antibody (Abcam, Alexa Fluor® 647, ab150075) were applied. Following staining, slides were mounted using Vectashield® Vibrance Antifade Mounting Medium with 4′,6-diamidino-2-phenylindole (DAPI) from Vector Laboratories (Newark, CA, H-1800). DAPI is a fluorescent stain commonly used to label cell nuclei, allowing for visualization of cellular morphology and nuclear structure. Imaging of the stained slides was acquired using a Zeiss Axioscan 7 Microscope, an automated slide scanner capable of capturing high-resolution images across large tissue sections. For quantitative analysis of the stained liver sections, the HALO® Image Analysis platform developed by Indica Labs (Albuquerque, NM, USA) was utilized. HALO® is specialized software designed for quantitative tissue analysis in digital pathology. Additionally, the HALO AI™ add-on module, which consists of neural network algorithm-driven tissue and cell classification and segmentation tools, was applied. This module aids in the automated identification and classification of specific cell populations within tissue sections. Using HALO® and HALO AI™, triple-stained liver sections (Ki67, ASGR1, DAPI) were analyzed to obtain a quantitative assessment of double positive Ki67+/ASGR1+ proliferating hepatocytes.

#### Mouse gene expression analysis

RNA from mouse tissues was extracted using the MagMAX™ mirVana™ Total RNA Isolation Kit (Thermo Fisher Scientific, A27828). cDNA was produced using the high-Capacity cDNA Reverse Transcription Kit (Thermo Fisher Scientific, 43-688-14) or the SuperScript™ IV VILO™ Master Mix (Thermo Fisher Scientific, 11756050). Mouse mRNA expression was quantified using TaqMan® Fast Advanced Master Mix (Thermo Fisher Scientific, 4444963). Probes used for each gene were as follows: *Alpl* (Mm00475834_m1), *Asgr1* (Mm00437595_m1), *Asgr2* (Mm00431863_m1), *Axin2* (Mm00443610_ml), *Ccnd1* (Mm00432359_ml), *Cyp1a2* (Mm00487224_ml), *Mki67* (Mm01278617_ml), *Notum* (Mm01253273_m1), *Wnt2* (Mm00470018_m1), *Wnt4* (Mm01194003_m1), *Wnt5a* (Mm01183986_m1), *Wnt5b* (Mm01183986_m1), *Wnt6* (Mm00437353_m1), *Wnt7b* (Mm01301717_m1), *Wnt8a* (Mm00442108_g1), *Wnt9b* (Mm00457102_m1), and *Wnt11* (Mm00437327_g1). Values were normalized to expression of the Actin B (*Actb*) housekeeping gene using the Mm02619580_gl probe (Thermo Fisher Scientific, 4351368).

#### Statistical Analysis

Statistical analysis of the data was conducted using GraphPad Prism 8 software (GraphPad Software, San Diego, CA). The significance level was set at p < 0.05. Results are presented as mean ± standard error of the mean (SEM). Statistical significance is denoted as follows: * p < 0.05, ** p < 0.01, *** p < 0.001, **** p < 0.0001. For RNA sequencing (RNAseq) data analysis, preprocessing steps, including alignment and quantification of reads, were performed using the STAR (Spliced Transcripts Alignment to a Reference) and RSEM (RNA-Seq by Expectation Maximization) packages. Subsequently, the DESeq2 package from the R software environment was utilized for differential gene expression analysis. Fold change, p-values, and false discovery rates (FDR) were calculated by DESeq2 to identify differentially expressed genes between experimental conditions.

**Figure S1.**
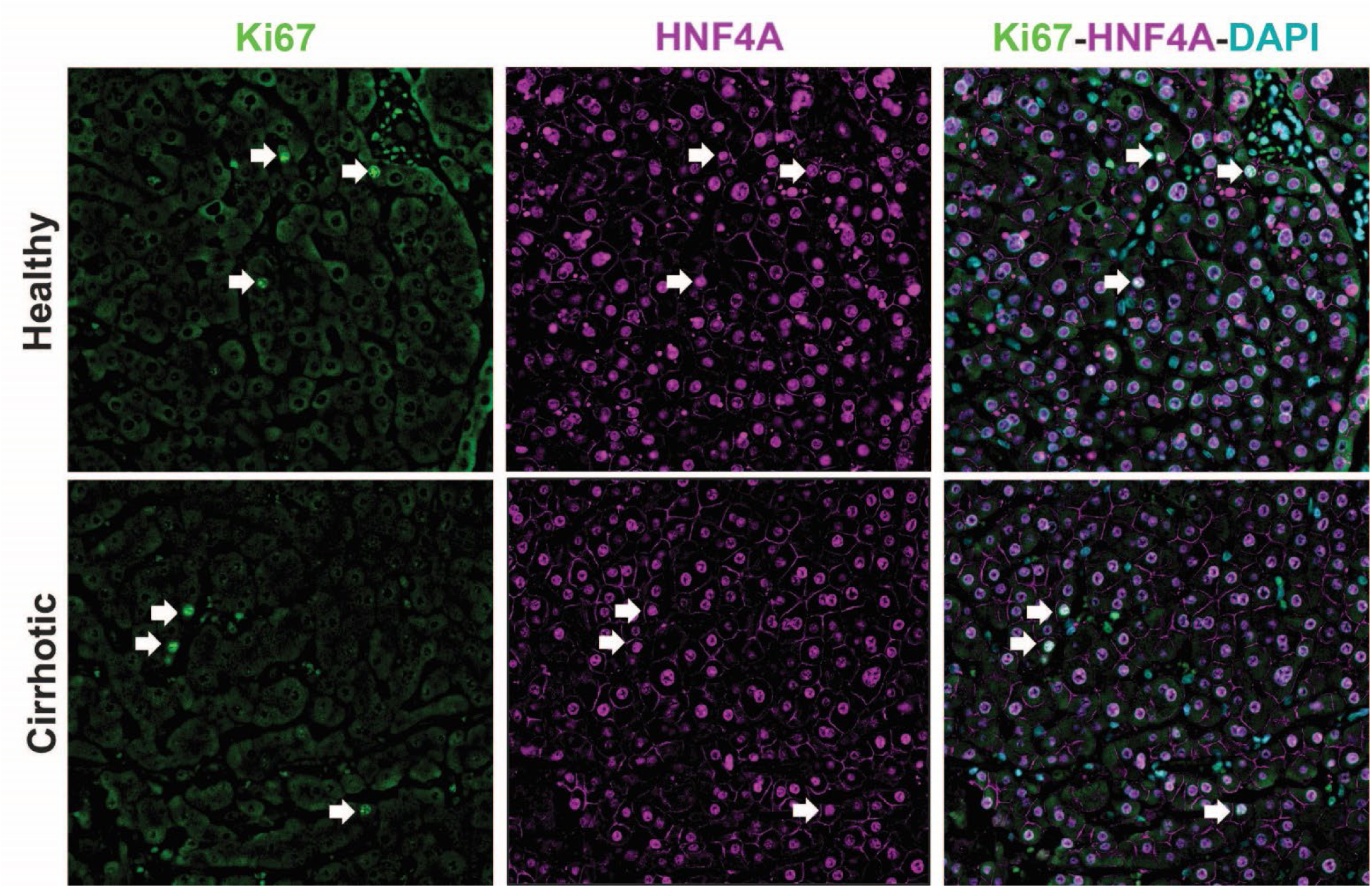
Proliferating hepatocytes in Alcohol-Associated Cirrhotic and Control Livers from Human Patients. Liver tissue sections were stained with the human hepatocyte specific, anti-human ASGR1 (red), and the proliferation marker, anti-Ki67 (green), antibodies, and DAPI (blue). White arrows show some hepatocytes staining for Ki67+ and HNF4A+. There were few proliferating hepatocytes in healthy and cirrhotic livers. There was no difference in the number of proliferating hepatocytes between both groups.

**Figure S2.**
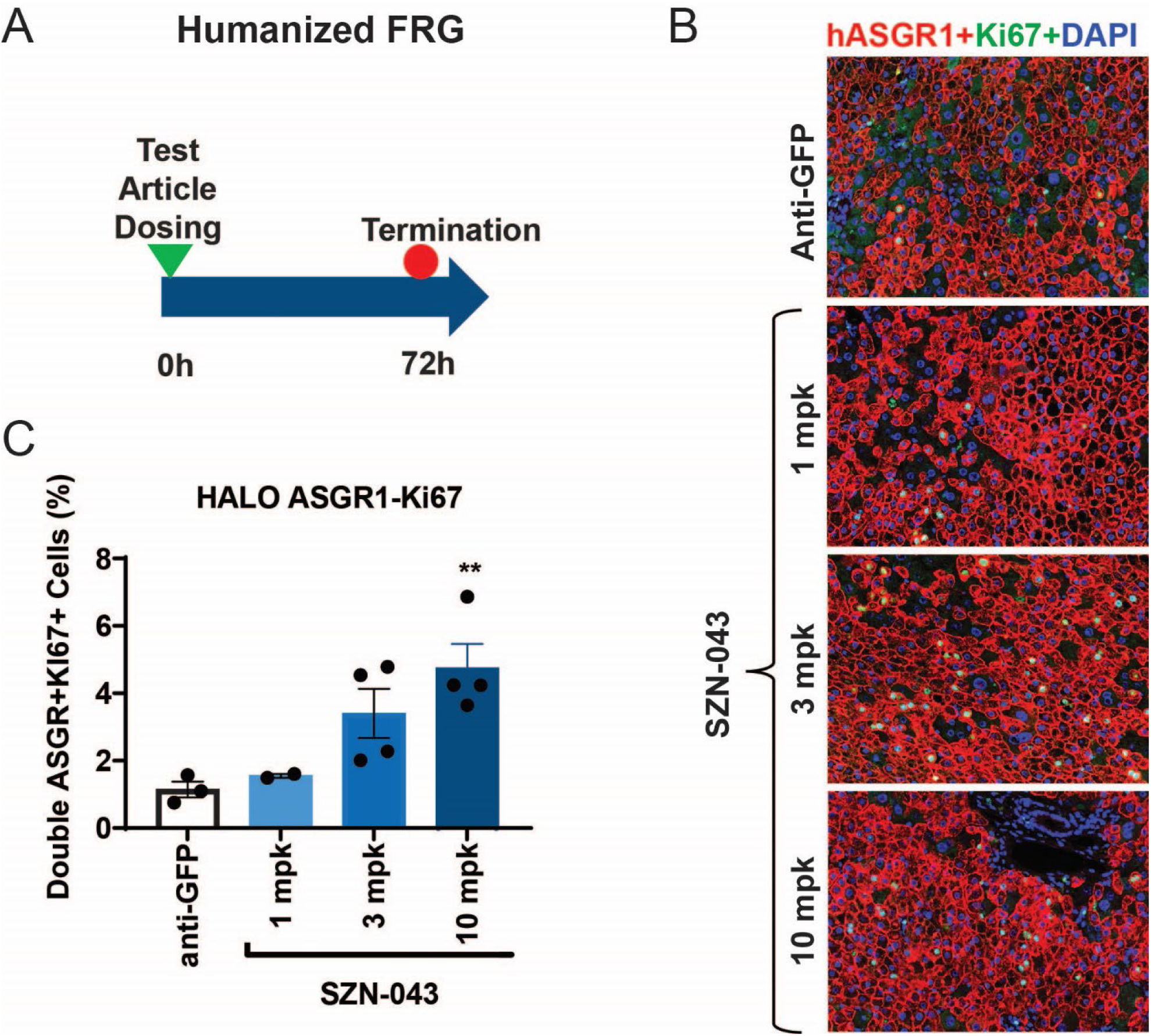
SZN-043 Dose-Dependent Induction of Proliferation in Human Hepatocytes. (A) Study design. FRG mice received a single dose, i.p., of either the negative control anti-GFP (10 mg/mL) or the SZN-043 test article at a 1, 3 or 10 mg/kg dose. The left lateral and medial liver lobes were collected upon termination on Day 3 for immunofluorescence analysis. (B) Left lateral liver tissue sections were stained with the human hepatocyte specific, anti-human ASGR1 (red), and the proliferation marker, anti-Ki67 (green), antibodies, and DAPI (blue) (C) The percentage of doubly positive cells was calculated using HALO imaging analysis software. ** p < 0.01

**Figure S3.**
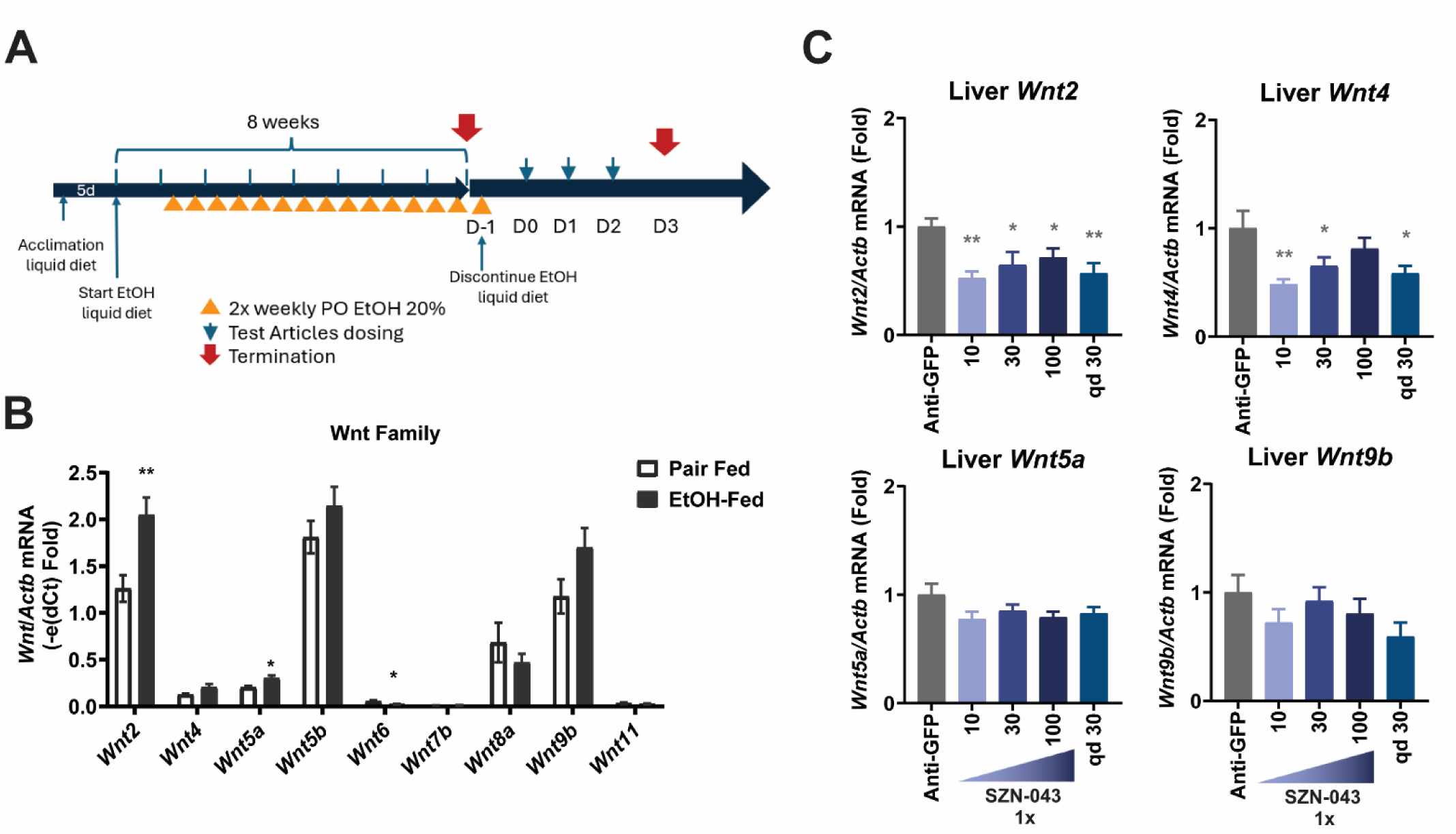
Effect of Ethanol and SZN-043 on the Expression of Wnt Family Members in Mice. (A) Study design. Aging C57BL/6J female mice were acclimated to the Lieber-DeCarli diet for 5 days, followed by an ethanol-supplemented liquid diet for 8 weeks. A control group was pair-fed with an equicaloric diet during this 8-week period. Starting on week 2 of the ethanol-supplemented diet, mice received 20% ethanol by oral gavage twice weekly for the remaining 7 weeks. The pair fed control group received an equivalent volume of ethanol-free equicaloric dose by oral gavage twice weekly also. Ethanol administration was then discontinued, and mice were returned to an ethanol-free liquid diet. The pair-fed control group and an ethanol-fed were euthanized upon ethanol discontinuation, on Day 0, and liver tissue was collected for analysis. 24 hours later, mice received either a single dose of SZN-043 at 10, 30 or 100 mg/kg, or daily dosing (30 mg/kg), or a single dose of a control antibody anti-GFP (30 mg/kg). On Day3, upon termination, liver tissue was collected for analysis. (B) QPCR expression analysis of Wnt family members in ethanol-treated or pair-fed controls. Results are presented as (e(dCt)) to reflect the relative abundance of different family members. (C) Effect of SZN-043 on *Wnt2*, *Wnt4*, *Wnt5a* and *Wnt9b* genes expression, determined by qPCR analysis.

**Figure S4.**
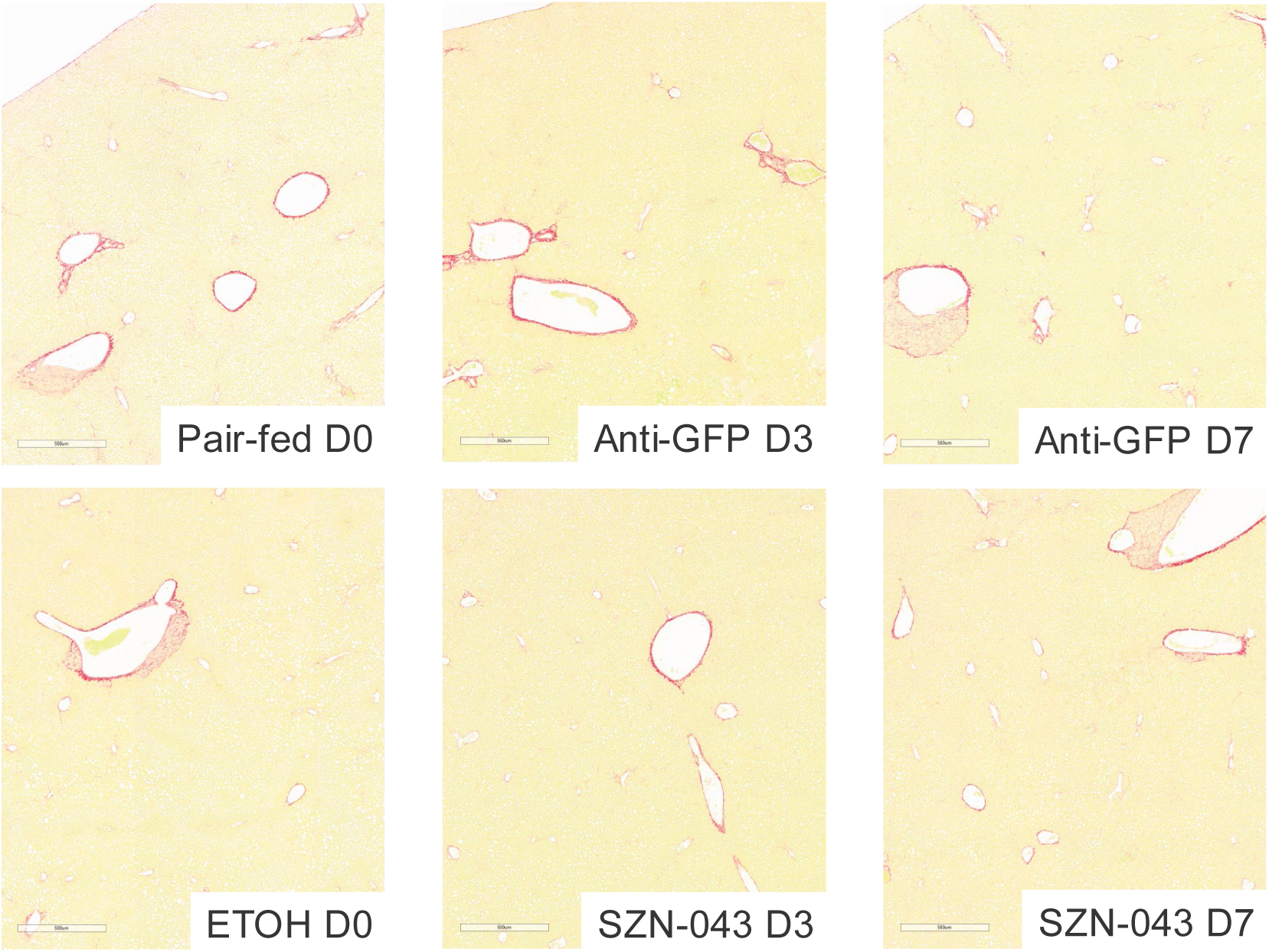
Picrosirius Red staining of Liver Sections from an Ethanol-Induced Injury Study. Study design. Aging C57BL/6J female mice were acclimated to the Lieber-DeCarli diet for 5 days, followed by an ethanol-supplemented liquid diet for 8 weeks. A control group was pair-fed with an equicaloric diet during this 8-week period. Starting on week 2 of the ethanol-supplemented diet, mice received 20% ethanol by oral gavage twice weekly for the remaining 7 weeks. The pair fed control group received an equivalent volume of ethanol-free equicaloric dose by oral gavage twice weekly also. Ethanol administration was then discontinued, and mice were returned to an ethanol-free liquid diet. The pair-fed control group and an ethanol-fed were euthanized upon ethanol discontinuation, on Day 0, and liver tissue was collected for analysis. Two hours later and daily thereafter until Day 6, mice received either SZN-043 (30 mg/kg) or a control antibody anti-GFP (30 mg/kg). On Day3 and on Day 7, upon termination, liver tissue was collected for analysis.

**Table S1.** Description of Human Healthy and Cirrhotic Liver Samples Used for Expression Analysis in Cohort #1.

| Sample | Gender | Age | Diagnostic | MELD |
| --- | --- | --- | --- | --- |
| **Control** |  |  |  |  |
| CT_01 | M | 68 | Malignant neoplasm of intrahepatic biliary conducts | n.a. |
| CT_02 | F | 63 | Benign hepatic neoplasm. Hepatic hemangioma. | n.a. |
| CT_03 | F | 55 | Secondary malignant hepatic neoplasm* | n.a. |
| CT_04 | M | 78 | Secondary malignant hepatic neoplasm | n.a. |
| CT_05 | M | 56 | Secondary malignant hepatic neoplasm* | n.a. |
| CT_06 | M | 80 | Secondary malignant hepatic neoplasm* | n.a. |
| CT_07 | F | 69 | Secondary malignant hepatic neoplasm* | n.a. |
| CT_08 | F | 70 | Secondary malignant hepatic neoplasm* | n.a. |
| CT_09 | F | 69 | Benign hepatic neoplasm | n.a. |
| CT_11 | M | 74 | Secondary malignant hepatic neoplasm* | n.a. |
| CT_12 | F | 64 | Secondary malignant hepatic neoplasm* | n.a. |
| CT_13 | M | 51 | Pancreatic neuroendocrine tumor | n.a. |
| **Cirrhotic** |  |  |  |  |
| CH_01 | M | 63 | Alcoholic cirrhosis | 25 |
| CH_02 | M | 48 | Alcoholic cirrhosis | 28 |
| CH_03 | M | 57 | Alcoholic cirrhosis | 13 |
| CH_04 | M | 58 | Alcoholic cirrhosis | 26 |
| CH_05 | M | 52 | Alcoholic cirrhosis | 19 |
| CH_06 | M | 67 | Alcoholic cirrhosis | 10 |
| CH_07 | M | 61 | Alcoholic cirrhosis | 13 |
| CH_09 | M | 65 | Alcoholic cirrhosis | 16 |
| CH_10 | M | 62 | Alcoholic cirrhosis | 14 |
| CH_11 | M | 64 | Alcoholic cirrhosis | 23 |
| CH_12 | M | 59 | Alcoholic cirrhosis | 20 |
| CH_13 | F | 56 | Alcoholic cirrhosis | 26 |

\* colorectal cancer metastasis

**Table S2.** Description of Healthy and Cirrhotic Liver Samples Used for Expression Analysis in Cohort #2.

| **Sample** | **Gender** | **Age** | **Diagnosis** | **MELD** |
| --- | --- | --- | --- | --- |
| HH1 | M | 70 | Control | n.a |
| HH3 | M | 59 | Control | n.a |
| HH4 | M | 54 | Control | n.a |
| HH5 | M | 26 | Control | n.a |
| HH6 | M | 50 | Control | n.a |
| ALC1 | M | 62 | Alcohol-associated cirrhosis | 16 |
| ALC2 | M | 43 | Alcohol-associated cirrhosis | 16 |
| ALC3 | F | 52 | Alcohol-associated cirrhosis | 15 |
| ALC5 | M | 45 | Alcohol-associated cirrhosis | 13 |

**Table S3.**
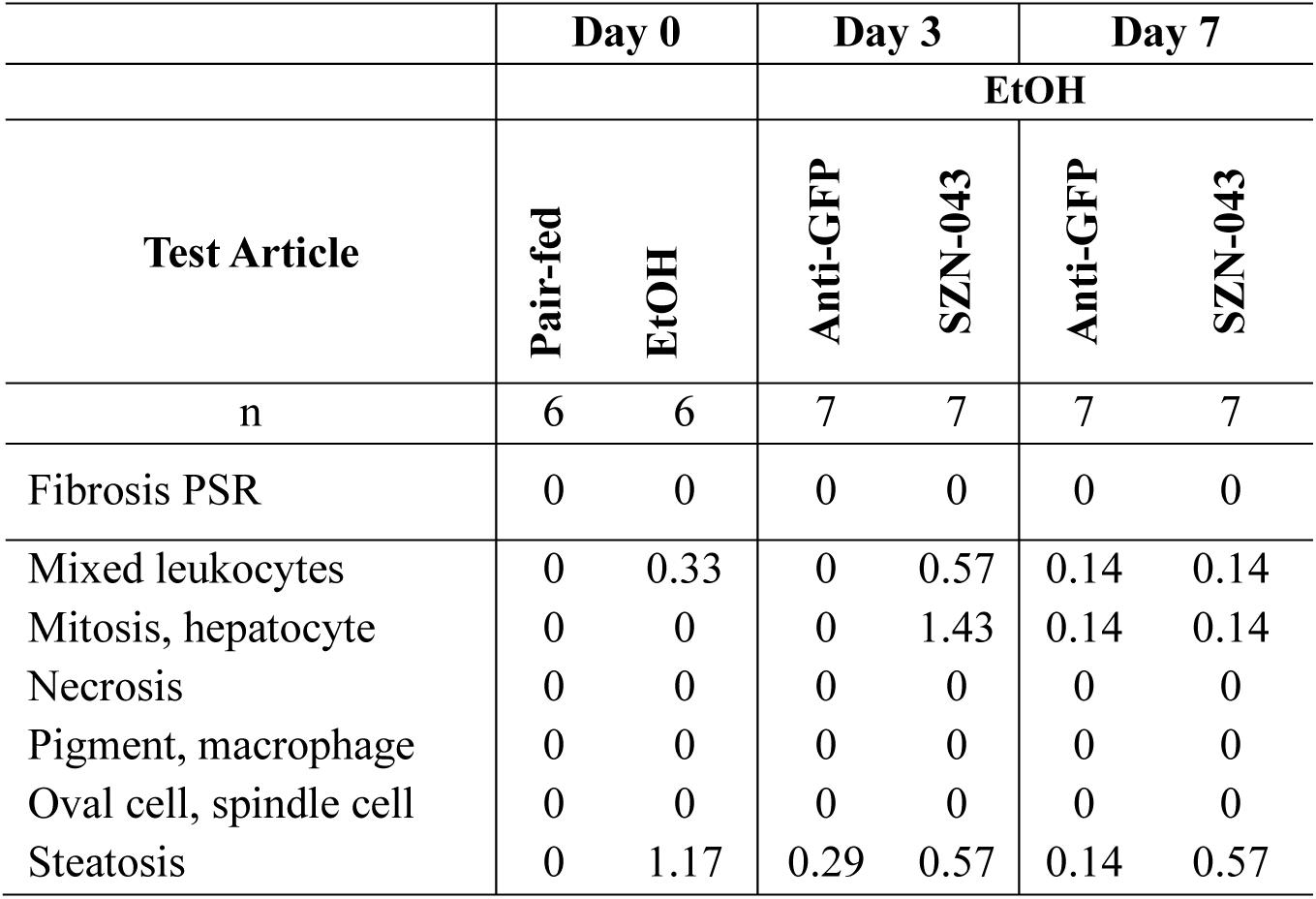
Summary of histological findings in ethanol-induced liver injury model.

## REFERENCES

1. Lee E, Navadurong H, Liangpunsakul S. Epidemiology and trends of alcohol use disorder and alcohol-associated liver disease. Clin Liver Dis (Hoboken) 2023;22:99–102.

2. Crabb DW, Im GY, Szabo G, Mellinger JL, Lucey MR. Diagnosis and Treatment of Alcohol-Associated Liver Diseases: 2019 Practice Guidance From the American Association for the Study of Liver Diseases. Hepatology 2020;71:306–333.

3. Singal AK, Bataller R, Ahn J, Kamath PS, Shah VH. ACG Clinical Guideline: Alcoholic Liver Disease. Am J Gastroenterol 2018;113:175–194.

4. Samala N, Gawrieh S, Tang Q, Lourens SG, Shah VH, Sanyal AJ, Liangpunsakul S, et al. Clinical Characteristics and Outcomes of Mild to Moderate Alcoholic Hepatitis. GastroHep 2019;1:161–165.

5. Lourens S, Sunjaya DB, Singal A, Liangpunsakul S, Puri P, Sanyal A, Ren X, et al. Acute Alcoholic Hepatitis: Natural History and Predictors of Mortality Using a Multicenter Prospective Study. Mayo Clin Proc Innov Qual Outcomes 2017;1:37–48.

6. Thursz MR. Screening and treatment of hepatitis B virus to prevent liver cancer in Africa. Hepat Oncol 2015;2:105–109.

7. Jindal A, Jagdish RK, Kumar A. Hepatic Regeneration in Cirrhosis. J Clin Exp Hepatol 2022;12:603–616.

8. Dubuquoy L, Louvet A, Lassailly G, Truant S, Boleslawski E, Artru F, Maggiotto F, et al. Progenitor cell expansion and impaired hepatocyte regeneration in explanted livers from alcoholic hepatitis. Gut 2015;64:1949–1960.

9. Lanthier N, Rubbia-Brandt L, Lin-Marq N, Clement S, Frossard JL, Goossens N, Hadengue A, et al. Hepatic cell proliferation plays a pivotal role in the prognosis of alcoholic hepatitis. J Hepatol 2015;63:609–621.

10. Yang Z, Zhang T, Kusumanchi P, Tang Q, Sun Z, Radaeva S, Peiffer B, et al. Transcriptomic Analysis Reveals the MicroRNAs Responsible for Liver Regeneration Associated With Mortality in Alcohol-Associated Hepatitis. Hepatology 2021;74:2436–2451.

11. Abu Rmilah A, Zhou W, Nelson E, Lin L, Amiot B, Nyberg SL. Understanding the marvels behind liver regeneration. Wiley Interdiscip Rev Dev Biol 2019;8:e340.

12. Russell JO, Monga SP. Wnt/beta-Catenin Signaling in Liver Development, Homeostasis, and Pathobiology. Annu Rev Pathol 2018;13:351–378.

13. Apte U, Singh S, Zeng G, Cieply B, Virji MA, Wu T, Monga SP. Beta-catenin activation promotes liver regeneration after acetaminophen-induced injury. Am J Pathol 2009;175:1056–1065.

14. Huang CK, Yu T, de la Monte SM, Wands JR, Derdak Z, Kim M. Restoration of Wnt/beta-catenin signaling attenuates alcoholic liver disease progression in a rat model. J Hepatol 2015;63:191–198.

15. Kaidi A, Williams AC, Paraskeva C. Interaction between beta-catenin and HIF-1 promotes cellular adaptation to hypoxia. Nat Cell Biol 2007;9:210–217.

16. Lehwald N, Tao GZ, Jang KY, Sorkin M, Knoefel WT, Sylvester KG. Wnt-beta-catenin signaling protects against hepatic ischemia and reperfusion injury in mice. Gastroenterology 2011;141:707–718, 718 e701-705.

17. Mao SA, Glorioso JM, Nyberg SL. Liver regeneration. Transl Res 2014;163:352–362.

18. Michalopoulos GK. Liver regeneration. J Cell Physiol 2007;213:286–300.

19. Monga SP, Pediaditakis P, Mule K, Stolz DB, Michalopoulos GK. Changes in WNT/beta-catenin pathway during regulated growth in rat liver regeneration. Hepatology 2001;33:1098–1109.

20. Benhamouche S, Decaens T, Godard C, Chambrey R, Rickman DS, Moinard C, Vasseur-Cognet M, et al. Apc tumor suppressor gene is the “zonation-keeper” of mouse liver. Dev Cell 2006;10:759–770.

21. Sekine S, Lan BY, Bedolli M, Feng S, Hebrok M. Liver-specific loss of beta-catenin blocks glutamine synthesis pathway activity and cytochrome p450 expression in mice. Hepatology 2006;43:817–825.

22. Soto-Gutierrez A, Gough A, Vernetti LA, Taylor DL, Monga SP. Pre-clinical and clinical investigations of metabolic zonation in liver diseases: The potential of microphysiology systems. Exp Biol Med (Maywood) 2017;242:1605–1616.

23. Lin Y, Fang ZP, Liu HJ, Wang LJ, Cheng Z, Tang N, Li T, et al. HGF/R-spondin1 rescues liver dysfunction through the induction of Lgr5(+) liver stem cells. Nat Commun 2017;8:1175.

24. Planas-Paz L, Orsini V, Boulter L, Calabrese D, Pikiolek M, Nigsch F, Xie Y, et al. The RSPO-LGR4/5-ZNRF3/RNF43 module controls liver zonation and size. Nat Cell Biol 2016;18:467–479.

25. Rocha AS, Vidal V, Mertz M, Kendall TJ, Charlet A, Okamoto H, Schedl A. The Angiocrine Factor Rspondin3 Is a Key Determinant of Liver Zonation. Cell Rep 2015;13:1757–1764.

26. Zebisch M, Xu Y, Krastev C, MacDonald BT, Chen M, Gilbert RJ, He X, et al. Structural and molecular basis of ZNRF3/RNF43 transmembrane ubiquitin ligase inhibition by the Wnt agonist R-spondin. Nat Commun 2013;4:2787.

27. Kim KA, Kakitani M, Zhao J, Oshima T, Tang T, Binnerts M, Liu Y, et al. Mitogenic influence of human R-spondin1 on the intestinal epithelium. Science 2005;309:1256–1259.

28. Zhang Z, Broderick C, Nishimoto M, Yamaguchi T, Lee SJ, Zhang H, Chen H, et al. Tissue-targeted R-spondin mimetics for liver regeneration. Sci Rep 2020;10:13951.

29. Nishikawa K, Osawa Y, Kimura K. Wnt/beta-Catenin Signaling as a Potential Target for the Treatment of Liver Cirrhosis Using Antifibrotic Drugs. Int J Mol Sci 2018;19.

30. Altamirano J, Miquel R, Katoonizadeh A, Abraldes JG, Duarte-Rojo A, Louvet A, Augustin S, et al. A histologic scoring system for prognosis of patients with alcoholic hepatitis. Gastroenterology 2014;146:1231–1239 e1231-1236.

31. Lackner C, Tiniakos D. Fibrosis and alcohol-related liver disease. J Hepatol 2019;70:294–304.

32. Hu S, Liu S, Bian Y, Poddar M, Singh S, Cao C, McGaughey J, et al. Single-cell spatial transcriptomics reveals a dynamic control of metabolic zonation and liver regeneration by endothelial cell Wnt2 and Wnt9b. Cell Rep Med 2022;3:100754.

33. Hu S, Monga SP. Wnt/-Catenin Signaling and Liver Regeneration: Circuit, Biology, and Opportunities. Gene Expr 2021;20:189–199.

34. Caldez MJ, Bjorklund M, Kaldis P. Cell cycle regulation in NAFLD: when imbalanced metabolism limits cell division. Hepatol Int 2020;14:463–474.

35. Grolmusz VK, Karaszi K, Micsik T, Toth EA, Meszaros K, Karvaly G, Barna G, et al. Cell cycle dependent RRM2 may serve as proliferation marker and pharmaceutical target in adrenocortical cancer. Am J Cancer Res 2016;6:2041–2053.

36. Cui H, Zhu X, Li S, Wang P, Fang J. Liver-Targeted Delivery of Oligonucleotides with N-Acetylgalactosamine Conjugation. ACS Omega 2021;6:16259–16265.

37. Bon C, Hofer T, Bousquet-Melou A, Davies MR, Krippendorff BF. Capacity limits of asialoglycoprotein receptor-mediated liver targeting. MAbs 2017;9:1360–1369.

38. Seitz HK, Xu Y, Simanowski UA, Osswald B. Effect of age and gender on in vivo ethanol elimination, hepatic alcohol dehydrogenase activity, and NAD+ availability in F344 rats. Res Exp Med (Berl) 1992;192:205–212.

39. Ramirez T, Li YM, Yin S, Xu MJ, Feng D, Zhou Z, Zang M, et al. Aging aggravates alcoholic liver injury and fibrosis in mice by downregulating sirtuin 1 expression. J Hepatol 2017;66:601–609.

40. Thursz MR, Richardson P, Allison M, Austin A, Bowers M, Day CP, Downs N, et al. Prednisolone or pentoxifylline for alcoholic hepatitis. N Engl J Med 2015;372:1619–1628.

41. Boulter L, Govaere O, Bird TG, Radulescu S, Ramachandran P, Pellicoro A, Ridgway RA, et al. Macrophage-derived Wnt opposes Notch signaling to specify hepatic progenitor cell fate in chronic liver disease. Nat Med 2012;18:572–579.

42. Louvet A, Thursz MR, Kim DJ, Labreuche J, Atkinson SR, Sidhu SS, O’Grady JG, et al. Corticosteroids Reduce Risk of Death Within 28 Days for Patients With Severe Alcoholic Hepatitis, Compared With Pentoxifylline or Placebo-a Meta-analysis of Individual Data From Controlled Trials. Gastroenterology 2018;155:458–468 e458.

43. Kim A, Wu X, Allende DS, Nagy LE. Gene Deconvolution Reveals Aberrant Liver Regeneration and Immune Cell Infiltration in Alcohol-Associated Hepatitis. Hepatology 2021;74:987–1002.

44. Niehrs C, Seidl C, Lee H. An “R-spondin code” for multimodal signaling ON-OFF states. Bioessays 2024;46:e2400144.

45. Tozawa R, Ishibashi S, Osuga J, Yamamoto K, Yagyu H, Ohashi K, Tamura Y, et al. Asialoglycoprotein receptor deficiency in mice lacking the major receptor subunit. Its obligate requirement for the stable expression of oligomeric receptor. J Biol Chem 2001;276:12624–12628.

46. Nioi P, Sigurdsson A, Thorleifsson G, Helgason H, Agustsdottir AB, Norddahl GL, Helgadottir A, et al. Variant ASGR1 Associated with a Reduced Risk of Coronary Artery Disease. N Engl J Med 2016;374:2131–2141.

47. Zhao W, Xu S, Weng J. ASGR1: an emerging therapeutic target in hypercholesterolemia. Signal Transduct Target Ther 2023;8:43.

48. Ueno S, Mojic M, Ohashi Y, Higashi N, Hayakawa Y, Irimura T. Asialoglycoprotein receptor promotes cancer metastasis by activating the EGFR-ERK pathway. Cancer Res 2011;71:6419–6427.

49. Zhu X, Song G, Zhang S, Chen J, Hu X, Zhu H, Jia X, et al. Asialoglycoprotein Receptor 1 Functions as a Tumor Suppressor in Liver Cancer via Inhibition of STAT3. Cancer Res 2022;82:3987–4000.

50. Kaffe E, Roulis M, Zhao J, Qu R, Sefik E, Mirza H, Zhou J, et al. Humanized mouse liver reveals endothelial control of essential hepatic metabolic functions. Cell 2023;186:3793–3809 e3726.

51. Hwang S, Ren T, Gao B. Obesity and binge alcohol intake are deadly combination to induce steatohepatitis: A model of high-fat diet and binge ethanol intake. Clin Mol Hepatol 2020;26:586–594.

52. Schonfeld M, O’Neil M, Villar MT, Artigues A, Averilla J, Gunewardena S, Weinman SA, et al. A Western diet with alcohol in drinking water recapitulates features of alcohol-associated liver disease in mice. Alcohol Clin Exp Res 2021;45:1980–1993.

53. Nautiyal N, Maheshwari D, Tripathi DM, Kumar D, Kumari R, Gupta S, Sharma S, et al. Establishment of a murine model of acute-on-chronic liver failure with multi-organ dysfunction. Hepatol Int 2021;15:1389–1401.

54. Takata N, Ishii KA, Takayama H, Nagashimada M, Kamoshita K, Tanaka T, Kikuchi A, et al. LECT2 as a hepatokine links liver steatosis to inflammation via activating tissue macrophages in NASH. Sci Rep 2021;11:555.

55. Tanida R, Goto H, Takayama H, Nakano Y, Oo HK, Galicia-Medina CM, Takahashi K, et al. LECT2 Deletion Exacerbates Liver Steatosis and Macrophage Infiltration in a Male Mouse Model of LPS-mediated NASH. Endocrinology 2024;165.

56. Wang JN, Li L, Li LY, Yan Q, Li J, Xu T. Emerging role and therapeutic implication of Wnt signaling pathways in liver fibrosis. Gene 2018;674:57–69.

57. Friedman SL, Pinzani M. Hepatic fibrosis 2022: Unmet needs and a blueprint for the future. Hepatology 2022;75:473–488.

58. Tsuchida T, Friedman SL. Mechanisms of hepatic stellate cell activation. Nat Rev Gastroenterol Hepatol 2017;14:397–411.

59. Bendell Jea. <Ph-1ab-OMP-131R10-anti-RSPO3-adv-solid-tumor-prev-treat-met-CRC-2016-EORTC-NCI-AACR.pdf>. 2017.

60. Nilsson KH, Henning P, El Shahawy M, Nethander M, Andersen TL, Ejersted C, Wu J, et al. RSPO3 is important for trabecular bone and fracture risk in mice and humans. Nat Commun 2021;12:4923.

61. Jacobsen FW, Stevenson R, Li C, Salimi-Moosavi H, Liu L, Wen J, Luo Q, et al. Engineering an IgG Scaffold Lacking Effector Function with Optimized Developability. J Biol Chem 2017;292:1865–1875.

62. Diehl KH, Hull R, Morton D, Pfister R, Rabemampianina Y, Smith D, Vidal JM, et al. A good practice guide to the administration of substances and removal of blood, including routes and volumes. J Appl Toxicol 2001;21:15–23.

63. Bertola A, Mathews S, Ki SH, Wang H, Gao B. Mouse model of chronic and binge ethanol feeding (the NIAAA model). Nat Protoc 2013;8:627–637.

